# Characterization of Immunoglobulin Loci in *Pongo abelii* and *Pongo pygmaeus*: Insights from Multi-Genome Annotation and IMGT-Based Curation

**DOI:** 10.1101/2025.10.22.683890

**Authors:** Shamsa Batool, Guilhem Zeitoun, Chahrazed Debbagh, Géraldine Folch, Joumana Jabado-Michaloud, Véronique Giudicelli, Sofia Kossida

**Affiliations:** IMGT®, IGH, Univ Montpellier, CNRS, Montpellier, France; Institut Universitaire de France (IUF), Paris, France

**Keywords:** IMGT, immunoinformatics, immunogenetics, immunoglobulins, IGH locus, IGL locus, IGK locus, *Pongo abelii*, *Pongo pygmaeus*

## Abstract

Antibodies, or immunoglobulins (IG), are central to the vertebrate adaptive immune system, yet the genomic architecture of IG loci remains poorly characterized in many nonhuman primates. In this study we present the first comprehensive genomic analysis of the immunoglobulin (IG) loci (IGH, IGL, and IGK) in two critically endangered orangutan species; *Pongo abelii* (Sumatran orangutan) and *Pongo pygmaeus* (Bornean orangutan) across multiple genome assemblies. Using IMGT-standardized biocuration framework combined with read-level structural validation, we identified previously undocumented haplotype-specific variation, including multigene duplications, asymmetric gene absence, and species-specific expansions of variable gene families. Recombination signal sequence (RSS) and switch region analyses revealed conserved regulatory motifs with potential implications for V(D)J recombination and class-switch recombination. These findings underscore the complexity and evolutionary adaptability of IG loci in great apes and highlight the value of orangutans as key references for understanding immune system evolution in the Hominidae lineage.

## 1. Introduction

Antibodies or immunoglobulins (IG) are the main molecules mediating humoral immune response. These molecules are present in all gnathostomes (jawed vertebrates), and are designed to neutralize pathogens by blocking their access to cells they might infect or destroy (Janeway, 2001). They comprise four polypeptide chains: two identical heavy (H) chains and two identical light immunoglobulin chains, which may be lambda (λ) or kappa (κ). The N-terminal region or Fab (fragment antigen-binding) is extremely diverse and responsible for recognizing and interacting with the antigen, while the C-terminal region or Fc (fragment crystallizable, the “antibody tail”) is responsible for triggering the effector cell responses. In placental mammals (eutheria), five classes of immunoglobulins are known to exist: IGHM, IGHD, IGHG, IGHE, and IGHA, each encoding a different class of IG (Das et al., 2012; Senger et al., 2015). They perform various effector functions, such as complement binding, binding to phagocytic cells, opsonization, and transport across the placental epithelium (Schroeder & Cavacini, 2010). Antibodies are not directly encoded in the germline genome; instead, they are produced through somatic V(D)J recombination processes, a site-specific recombination process that joins variable (V), diversity (D), and joining (J) genes to generate antibody diversity (Tonegawa, 1983). V(D)J rearrangements are facilitated by the recombination activating gene (RAG) proteins, RAG-1 and RAG-2. These proteins are responsible for recognizing and cleaving DNA at specific sites, producing precise double-strand breaks at the junctions between the recombination signal sequences (RSSs) and the coding regions (Gellert, 1997a). This process affects the immunoglobulin (IG) loci that contain variable (V), diversity (D), and joining (J) genes. In the heavy chain, recombination occurs in two steps: first, the D and J genes are joined, followed by the addition of a V gene to form the complete V(D)J exon that encodes the variable region of the antibody. In contrast, light chains (kappa or lambda) lack D genes and are assembled by recombining only V and J genes. The antibodies are then diversified by somatic hypermutation, which introduces point mutations in certain hotspots (Dudley et al., 2005). Immunoglobulin loci have diversified rapidly and independently across jawed vertebrate lineages, resulting in major differences in their genomic organization, gene content and the proportion of functional genes versus pseudogenes, reflecting strong lineage-specific evolutionary pressures (Das et al., 2012).

IMGT®, the international ImMunoGeneTics information system® (Manso et al., 2022), was created in 1989. The establishment of IMGT® marked the birth of immunoinformatics—a new scientific field emerging at the intersection of immunogenetics and bioinformatics. For the first time, immunoglobulin (IG or antibody) and T cell receptor (TR) variable (V), diversity (D), joining (J), and constant (C) genes were officially acknowledged as true “genes,” as well as the conventional genes (Lefranc & Lefranc, 2001a, 2001b). This significant breakthrough enabled the organization and management of genes and data related to the complex and highly diverse adaptive immune responses within genomic databases and tools using standardized IMGT nomenclature.

Nonhuman primates are of great interest in comparative studies and biomedical research due to their close similarities with humans (Grow et al., 2016). Several studies have explored the relationship between nonhuman primate evolution and human disease, with a particular focus on segmental duplications observed in great apes and humans, which are thought to play an important role in human disease susceptibility, as in the case of their impact on genes associated with Mendelian diseases (Bailey & Eichler, 2006; Symmons et al., 2008). Studies of adaptive immune responses across various vertebrate species open new therapeutic opportunities (Muyldermans & Smider, 2016). Gaining insight into the diversity of adaptive immune systems across different vertebrates can also aid in examining the transmission and spread of novel zoonotic pathogens (Bean et al., 2013; Roffler et al., 2024).

Orangutans were once classified into two subspecies. However, recent research and taxonomic revisions—driven by extensive molecular and morphological analyses have shown that the genetic and physical differences between Sumatran and Bornean orangutans are as significant as, or even greater than, those between the two chimpanzee species, the common chimpanzee and the bonobo. (Ferris et al., 1981; Janczewski et al., 1990; Ruvolo et al., 1994; Ryder & Chemnick, 1993; Warren et al., 2001; Xu & Arnason, 1996; Zhi et al., 1996). As a result, updated classifications now recognize Sumatran and Bornean orangutans as distinct species; *Pongo abelii* in Sumatra and *Pongo pygmaeus* in Borneo, with an estimated divergence time of around 1.1 million years ago (Groves, 2001; Warren et al., 2001). However, dissenting opinions exist (Muir et al., 1998), noting that despite a pericentric inversion in chromosome 2, interbreeding between these groups in captivity can still produce fertile offspring (Seuanez et al., 1979). A third species, *Pongo tapanuliensis*, identified in the Batang Toru region of Sumatra, diverged from *Pongo abelii* around 3.4 million years ago based on genomic and morphological data (Nater et al., 2017). Because orangutans (genus *Pongo*)—the only Asian great apes are the most distantly related to humans within the Hominidae family, they hold particular significance in studies exploring evolutionary relationships among great apes. The orangutan genome was published in 2011 (Locke et al., 2011), revealing a strikingly slower rate of structural genome evolution in the *Pongo* lineage. Assuming consistent mutation rates and generation times across the Hominidae family, the most recent common ancestor of orangutans and the other great apes is estimated to have lived between 15 and 21 mya (Groves, 2001; Perelman et al., 2011; Prado-Martinez et al., 2013; Schrago & Voloch, 2013).

Our analysis of IG loci in two orangutan species (*Pongo abelii* and *Pongo pygmaeus*), aiming to identify both shared and distinct features between these two species and humans provides valuable insights into the evolution of immune response mechanisms and immune system adaptability across great apes. While great apes represent the most appropriate model for evaluating the safety and efficacy of vaccines and treatments intended for human use, concerns regarding endangered species status, cost and limited availability pose significant ethical and practical challenges and make the use of great apes, including orangutans, highly restricted (Aguilera et al., 2021). Nevertheless, genomic data from these species remain essential in experimental research for advancing our understanding of immunological diseases and supporting the development of potential therapies. This information can provide insights into the differences in primate immune systems and how they have evolved to respond to infections. Likewise, it should have implications for biomedical trials for drug development and evaluation (Bjornson-Hooper et al., 2022).

## 2. Materials and methods

### 2.1 Genomic dataset selection and preliminary assessment

To investigate the immunoglobulin (IG) loci in orangutans, we analyzed all available genome assemblies (n=13) as of January 2024 from the NCBI Genome database. These assemblies represented two orangutan species (*Pongo abelii* & *Pongo pygmaeus*) and 7 assemblies were selected using IMGT’s predefined two-phase pre-selection criteria (Papadaki et al., 2025).

### 2.2 Long read based assembly validation

Following selection, we used IMGT/StatAssembly (version v0.1.4) (Sanou et al., 2025; Zeitoun et al., 2025) as part of IMGT-developed quality control process incorporating long-read sequencing data retrieved from NCBI BioProjects. Preference was given to HiFi reads due to their medium range but high quality; in cases where HiFi data were unavailable, PacBio CCS or the highest-quality long reads were used. Raw reads underwent quality assessment using FastQC (Andrews, 2010), followed by locus-specific alignment using minimap2 (2.27-r1193) (Li, 2021) with the -x map-hifi preset. The BAM file was generated from reads based on the script (https://src.koda.cnrs.fr/imgt-igh/statassembly/-/blob/master/example_files/assembly.py?ref_type=heads). The pipeline computed key metrics, including coverage depth, breadth, mapping quality, mismatch and indel rates, and structural variant support. This workflow ensured high-confidence, structurally validated assemblies for downstream IG gene analysis.

### 2.3 Locus delimitation, extraction and integration in IMGT/LIGM-DB

Locus extraction, as previously described in the IMGT biocuration framework (Pégorier et al., 2020; Nguefack Ngoune et al., 2022; Papadaki et al., 2025) was initiated by comparing the assembly with the existing and most recent human IMGT reference directory sets, using BLAST (version BLAST+ 2.16.0), targeting the immunoglobulin heavy (IGH), kappa (IGK), and lambda (IGL) loci. For each locus, boundaries were identified based on the presence of 5’ and 3’ terminal IG genes, with an additional 10kb extension on both ends to capture potential unannotated genes. For IGH, IGK and IGL loci, locus delimitation relied on conserved flanking non-IG genes known as IMGT “bornes” (http://www.imgt.org/IMGTindex/IMGTborne.php) such as PAX8 and RPIA for IGK, TOP3B and RSPH14 for IGL and TMEM121 for IGH. For the remaining assemblies, locus extraction was guided by alignment to the first assembly annotations, which served as the reference. For loci that fulfilled all IMGT quality criteria, corresponding IMGT/LIGM-DB (Giudicelli et al., 2006) entries were created, with unique IMGT accession numbers.

### 2.4 Gene Annotation and Functionality Determination

Following extraction, IMGT/LIGMotif (Lane et al., 2010) was used to analyze, identify and classify V (variable), D (diversity), J (joining), and C (constant) genes and the outcome was integrated, visualized, and verified in VectorNTI (v. 11.5.3, Thermo Fisher). The annotation rules described in the IMGT scientific chart are based on the axioms of IMGT-ONTOLOGY (Giudicelli & Lefranc, 1999) and allow the detailed characterization of the gene and thus, its functionality definition (Functional, ORF, or Pseudogene) (https://www.imgt.org/IMGTindex/functionality.php) based on sequence integrity and comparison with reference prototypes (https://www.imgt.org/IMGTindex/prototypes.php). The coding regions for IG V genes were analyzed with IMGT/V-QUEST (Giudicelli et al., 2004) and validated through multiple alignment tools including Clustal Omega tool/EMBL-EBI (Madeira et al., 2019), to ensure framework regions (FR-IMGT) and complementarity determining region (CDR-IMGT) delimitation consistency across IG. Annotated genes and alleles were integrated into IMGT/LIGM-DB (Giudicelli et al., 2006), IMGT/GENE-DB (Giudicelli et al, 2004) and the IMGT reference directory (https://www.imgt.org/vquest/refseqh.html). All the annotated genomic data are available in the corresponding IMGT repertoire pages (Locus representation, Locus description, Locus in genome the assembly, Locus gene order, Locus Borne, Gene tables, Potential germline repertoire, Protein displays, Alignments of alleles, Colliers de Perles, and germline [CDR1-IMGT.CDR2-IMGT.CDR3-IMGT] lengths) and the IMGT reference directory available at https://www.imgt.org/IMGTrepertoire/ and https://www.imgt.org/vquest/refseqh.html, respectively.

The IMGT/LIGM-DB annotated locus sequences were submitted to the NCBI Third Party Annotation (TPA) (Cochrane et al., 2006) and TPA accession numbers were assigned. These numbers replace the original IMGT accession numbers, however, these latter will remain always valid for database requests (https://www.imgt.org/IMGTrepertoire/numacc.php). This dual referencing ensures full correspondence between IMGT records and publicly accessible NCBI entries, facilitating transparent cross-referencing and traceability of annotated immunoglobulin loci across databases.

### 2.5 IMGT Nomenclature

The nomenclature of *Pongo* immunoglobulin (IGH, IGK, and IGL) genes and alleles follows the rules of the standardized IMGT® nomenclature, (accessible via https://www.imgt.org/IMGTScientificChart/Nomenclature/IMGTnomenclature.php) (Lefranc, 2014). Gene and allele names convey specific information regarding the gene type, allelic variant, and genomic position within the locus. For variable (V) genes, including IGHV, IGKV, and IGLV, the names indicate the subgroup, a set of genes sharing at least 75% nucleotide identity (https://www.imgt.org/IMGTindex/subgroup.php) and relative position from the 3′ to 5′ end of the locus. Pseudogenes that cannot be assigned to a subgroup due to low % (>75%) of identity of the V-REGION (degenerated and/or truncated pseudogenes) are assigned to a Clan (https://www.imgt.org/IMGTindex/Clan.php). The name of the clan is indicated with a Roman number between parentheses based on homology with the closest species. For diversity (D) and joining (J) genes (IGHD, IGHJ, IGKJ and IGLJ), the nomenclature reflects the set and position from 5′ to 3′, with numbering increasing along the locus. Constant (C) genes are designated according to the encoded immunoglobulin isotype in the IGH, whereas in IGL, with multiple C genes, the numbering increases from 5′ to 3. When one or more newly identified genes are located between two already known and named genes within the same locus, an additional hyphen and number are added to indicate their positions in the insertion (e.g., IGKV2-55-1 between IGKV2-55 and IGKV2-56 in IGK locus of *Pongo abelii*).

An allele represents a polymorphic variant of a gene, defined by nucleotide-level differences within its core sequence (V-, D-, J-REGION, or C exons, depending on gene type). Alleles are designated by the gene name followed by an asterisk and a two-digit number starting from 01. New alleles were identified by comparing their coding regions (V-REGION for IGHV, IGKV, and IGLV; D-REGION for IGHD; J-REGION for IGHJ, IGKJ, and IGLJ; C-REGION for IGKC and IGLC; and exons for IGH constant genes) with the IMGT reference directory sets of the same species. The IMGT® reference directories contain the curated reference sequences for all IGH, IGK, and IGL gene alleles.

### 2.6 Validation of novel alleles and Structural variants

Given that the immunoglobulin (IG) loci are highly complex and polymorphic genomic regions enriched with segmental duplications and highly similar gene families, they are prone to misassemblies or collapsed regions in standard genome assemblies (Zhu et al., 2024). To resolve these complexities and to validate haplotype-specific duplications and insertions identified in the IG loci, we performed read-level analysis. Using IMGT/StatAssembly, BAM alignments of HiFi and Oxford Nanopore reads retrieved from the NCBI BioProjects PRJNA916742 and PRJNA916743 were used to assess read coverage and consistency across exonic regions. Regions containing genes present exclusively on one haplotype were closely examined to distinguish true structural variation from potential assembly artifacts. We aligned the raw Nanopore reads back to the corresponding assemblies and visualized these regions in IGV (Integrative Genomics Viewer v2.18.5+dfsg-1). For each candidate duplication or insertion, we confirmed the presence of continuous long read spanning the variant region, providing strong evidence for their authenticity. This approach ensured that both novel alleles and observed haplotype-specific gene gains were supported by underlying read data and not artifacts of genome assembly or misalignment.

Alleles and genes that failed to meet validation standards were excluded or retracted, with data updates tracked in IMGT/GENE-DB (available at https://www.imgt.org/IMGTgenedbdoc/dataupdates.html).

### 2.7 Analysis of RSS

For functional IG V, D, and J genes, the corresponding recombination signal sequences (RSSs) were extracted from the IMGT database for the IGH, IGK, and IGL loci to assess the degree of conservation and variation of recombination signal sequences (RSSs) in *Pongo*. These sequences include the conserved heptamer and nonamer motifs separated by a spacer typically 12 or 23 base pairs (bp) flanking the coding segments, which are essential for RAG-mediated V(D)J recombination (Gellert, 1997a). Sequence logos were generated in Python (Matplotlib v3.10.1) using position frequency matrices (PFMs) in a WebLogo-style layout (Crooks et al., 2004; Schneider & Stephens, 1990). Spacer length distributions were shown between the motif panels.

### 2.8 Identification and analysis of switch regions

To investigate the organization and conservation of potential immunoglobulin switch (S) regions in orangutans, genomic regions upstream of each constant region (IGHC) gene were examined for characteristic sequence features associated with class-switch recombination. Coordinates of IGHC genes were obtained from annotated genome assemblies of *Pongo abelii* and *Pongo pygmaeus* (GCF_028885655.2 & GCF_028885625.2 respectively). For each gene, a 5-kb sequence upstream of the transcription start site (TSS) was extracted from the indexed genome using BEDTools getfasta, with strand orientation taken into account. These upstream regions, which are expected to encompass the immunoglobulin switch (S) regions, were screened for degenerate pentamer motifs (GAGCW, where W = A or T; and GGGBT, where B = C, G, or T) representing canonical AID-target sequences (Chaudhuri & Alt, 2004; Mills et al., 1990). Motif counts and GC content were computed using custom Python scripts and normalized per kilobase to allow comparison among genes. Local repeat structures within each upstream region and across the entire IGHC cluster were visualized by self-dotplot analysis in standalone Gepard application (Krumsiek et al., 2007) using a word size of 14.

### 2.9 Comparative and Phylogenetic Analysis of the Immunoglobulin Loci Across Four Species

Immunoglobulin variable (V) gene clusters (V-CLUSTER) for the heavy (IGHV) and light chains (IGKV, IGLV) were analyzed across multiple genome assemblies of *Homo sapiens*, *Pongo abelii*, *Pongo pygmaeus*, and *Gorilla gorilla gorilla*. For each genome, the length of each V-CLUSTER (in base pairs) and the number of V genes were recorded. Gene density was calculated as the number of V genes per megabase (genes/Mb) of the cluster using the formula:

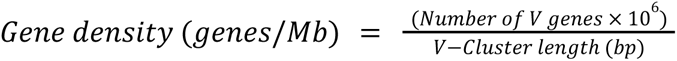

Densities were calculated separately for each genome assembly and then averaged across assemblies within each species to obtain mean values and standard deviations for each locus. Comparative analyses and visualizations were performed in Python using *pandas* and *matplotlib*. Locus size versus gene number relationships were visualized as scatterplots, and species-level summaries as bar charts showing mean density ± SD.

Phylogenetic relationships among immunoglobulin variable (V) genes from the IGHV, IGKV, and IGLV loci of *Homo sapiens, Pongo abelii, Pongo pygmaeus*, and *Gorilla gorilla gorilla* were analyzed using NGPhylogeny “One Click” online workflow (Lemoine et al., 2019). All annotated V genes were represented by their 01 alleles (IMGT reference sequences). The pipeline performs multiple sequence alignment with MUSCLE (Edgar, 2004) and constructs maximum-likelihood trees using PhyML (Guindon & Gascuel, 2003). Bootstrap support values (100 replicates) were calculated automatically, and resulting trees were visualized in iTOL (Letunic & Bork, 2021) using circular layouts. Branches were color-coded by species, and color strips indicated IGHV, IGKV, and IGLV subgroup classifications.

## 3. Results

### 3.1 Dataset selection

All genome assemblies used for the annotation of immunoglobulin (IG) loci were subjected to the IMGT quality control process (Supplementary Table 1). Each assembly underwent comprehensive assessment based on sequence accuracy, structural consistency, and read support, incorporating long-read sequencing data from NCBI BioProjects. Only assemblies that successfully passed IMGT’s quality thresholds were included for downstream IG locus annotation and comparative analysis. This ensured that the identified IGH, IGL and IGK loci were derived from high-confidence assemblies, minimizing artifacts related to sequencing errors or assembly discrepancies (Supplementary Figure 1 & 2). Table 1 provides a comprehensive summary of the information of the selected assemblies, including locus positions and corresponding accession numbers within the IMGT/LIGM-DB (v1.2.11).

**Table 1.** Summary of genome assemblies and immunoglobulin (IG) loci annotated in *Pongo* species. This table summarizes the annotated genome assemblies of *Pongo abelii* (Sumatran orangutan) and *Pongo pygmaeus* (Bornean orangutan) and the corresponding immunoglobulin heavy (IGH) and lambda light (IGL) chain loci identified within them. For each assembly, the GenBank accession identifiers, chromosomal or scaffold coordinates, and locus orientations are provided along with the associated IMGT/LIGM-DB accession numbers. * Chromosome numbers differ in the latest telomere-to-telomere (T2T) assemblies due to updated chromosome nomenclature and reannotation of genomic structure (https://ncbiinsights.ncbi.nlm.nih.gov/2024/05/13/refseq-release-224/). ** Reference assembly for *Pongo abelii* in IMGT databases

| Genome assembly | Isolate | GenBank assembly ID | Chromosome GenBank assembly ID /scaffold ID | Chromosome GenBank assembly/ Contig locus positions | Locus length (bp) | IMGT locus orientation on chromosome | IMGT/LIGM-DB /TPA accession number |
| --- | --- | --- | --- | --- | --- | --- | --- |
| IGH on chromosome 14/15* |  |  |  |  |  |  |  |
| *Pongo abelii* (Sumatran orangutan), NCBI taxon:9601 |  |  |  |  |  |  |  |
| Susie_PABv2** | Susie | GCA_002880775.3 | CM009276.2 | 87402090..88963417 | 1561328 | REVERSE | IMGT000122 |
| NHGRI_mPonAbe1-v2.1_pri | AG06213 | GCA_028885655.3 | NC_072000.2 | 110080644..111679704 | 1492363 | REVERSE | IMGT000206 |
| NHGRI_mPonAbe1-v2.0_alt | AG06213 | GCA_028885685.2 | CM054671.2 | 102190285..103682647 | 1599061 | REVERSE | IMGT000121 |
| Susie_PAB_hifiasm-v0.15.2.pri | Susie | GCA_030170355.1 | JARRAK010000025.1 | 89157563..90696389 | 1538827 | - | IMGT000157 |
| Susie_PAB_hifiasm-v0.15.2.alt | Susie | GCA_030170345.1 | JARRAL010000083.1 | 27984844..29453109 | 1468266 | - | IMGT000167 |
| *Pongo pygmaeus* (Bornean orangutan), NCBI taxon:9600 |  |  |  |  |  |  |  |
| NHGRI_mPonPyg2-v2.1_pri | AG05252 | GCA_028885625.3 | NC_072388.2 | 104726350..106399170 | 1745983 | REVERSE | IMGT000230 |
| NHGRI_mPonPyg2-v2.0_alt | AG05252 | GCA_028885525.2 | CM054622.2 | 105311893..107057879 | 1672821 | REVERSE | IMGT000229 |
| IGL locus on chromosome 22/23* |  |  |  |  |  |  |  |
| *Pongo abelii* (Sumatran orangutan), NCBI taxon:9601 |  |  |  |  |  |  |  |
| Susie_PABv2 | Susie | GCA_002880775.3 | CM009284.2 | 3326936-4190098 | 863163 | REVERSE | IMGT000108 |
| NHGRI_mPonAbe1-v2.1_pri | AG06213 | GCA_028885655.3 | NC_085929.1 | 25547691..26403226 | 855536 | REVERSE | IMGT000246 |
| NHGRI_mPonAbe1-v2.0_alt | AG06213 | GCA_028885685.2 | CM068846.1 | 21999085..22854931 | 855847 | REVERSE | IMGT000247 |
| Susie_PAB_hifiasm-v0.15.2.pri | Susie | GCA_030170355.1 | JARRAK010000033.1 | 17758621..18622348 | 150504 | - | IMGT000189 |
| Susie_PAB_hifiasm-v0.15.2.alt | Susie | GCA_030170345.1 | JARRAL010000044.1 | 31155160..32018390 | 950543 | - | IMGT000190 |
| *Pongo pygmaeus* (Bornean orangutan), NCBI taxon:9600 |  |  |  |  |  |  |  |
| NHGRI_mPonPyg2-v2.1_pri | AG05252 | GCA_028885625.3 | NC_085931.1 | 29525211..30907261 | 1382051 | REVERSE | IMGT000278 |
| NHGRI_mPonPyg2-v2.0_alt | AG05252 | GCA_028885525.2 | CM068910.1 | 20975388..22354260 | 1378873 | REVERSE | IMGT000248 |
| IGK locus on chromosome 2A/12* |  |  |  |  |  |  |  |
| *Pongo abelii* (Sumatran orangutan), NCBI taxon:9601 |  |  |  |  |  |  |  |
| Susie_PABv2 | Susie | GCA_002880775.3 | CM009263.2 | 19648734-20476838 | 828105 | FORWARD | IMGT000096<br>BK063599 |
| NHGRI_mPonAbe1-v2.1_pri | AG06213 | GCA_028885655.3 | NC_071997.2 | 43451819-44286661 | 834843 | FORWARD | IMGT000244 |
| NHGRI_mPonAbe1-v2.0_alt | AG06213 | GCA_028885685.2 | CM054668.2 | 42039675..42823442 | 783768 | FORWARD | IMGT000245 |
| Susie_PAB_hifiasm-v0.15.2.pri | Susie | GCA_030170355.1 | JARRAK010000016.1 | 80163254..80977396 | 814143 | - | IMGT000201 |
| Susie_PAB_hifiasm-v0.15.2.alt | Susie | GCA_030170345.1 | JARRAL010000033.1 | 15005270..15802831 | 797562 | - | IMGT000202 |
| *Pongo pygmaeus* (Bornean orangutan), NCBI taxon:9600 |  |  |  |  |  |  |  |
| NHGRI_mPonPyg2-v2.1_pri | AG05252 | GCA_028885625.3 | NC_072385.2 | 44675560..45472918 | 797359 | FORWARD | IMGT000275 |
| NHGRI_mPonPyg2-v2.0_alt | AG05252 | GCA_028885525.2 | CM054619.2 | 47958023..48790500 | 832478 | FORWARD | IMGT000276 |

#### 3.1.1 Susie

For the *Pongo abelii* individual Susie, we analyzed three available genome assemblies: Susie_PABv2 (GCA_002880775.3), Susie_PAB_hifiasm-v0.15.2.pri (GCA_030170355.1), and Susie_PAB_hifiasm-v0.15.2.alt (GCA_030170345.1). The Susie_PABv2 released in 2018 serves as the reference assembly in the IMGT database and represents a more complete, chromosome-scale sequence, while the latter two released in 2023 generated using the Hifiasm assembler correspond to the primary and alternate haplotypes, respectively, and are estimated to be more accurate (quality value [QV] = 42–58 or 99.9937%–99.9998% accuracy) and significantly more contiguous (contig N50 = 19–104 Mbp) (Mao et al., 2024). The Hifiasm-based assemblies do not fully meet IMGT quality criteria due to incomplete contiguity and lack of full scaffolding and assembly polishing Supplementary Table 1. Nevertheless, they offer valuable phased views of the IG loci, enabling detection of allelic variation, hemizygosity, and haplotype-specific structural events that are not readily captured in the haploid representation of Susie_PABv2. By analyzing and annotating IG genes across all three assemblies, we utilized the high continuity of the reference genome and the resolution of haplotype-specific variation achieved through long-read phasing to obtain a more complete understanding of IG diversity in this individual.

#### 3.1.2 Telomere to telomere (T2T) assemblies alter chromosomal localization of IG loci

We analyzed four telomere-to-telomere (T2T) assemblies released in 2024, NHGRI_mPonPyg2-v2.1_pri (GCF_028885625.2), NHGRI_mPonPyg2-v2.1_alt (GCA_028885525.2), NHGRI_mPonAbe1-v2.1_pri (GCF_028885655.2), and NHGRI_mPonAbe1-v2.1_alt (GCA_028885685.2) representing phased genome sequences for *Pongo pygmaeus* (AG05252) and *Pongo abelii* (AG06213). These assemblies employ a revised chromosome numbering convention based on alignment to a common great ape and human reference, resulting in reassigned chromosome identifiers better reflecting evolutionary and structural correspondence. For instance, the IGH locus, previously annotated on chromosome 14, is now located on chromosome 15; the IGK locus shifts from chromosome 2A to 12; and the IGL locus from 22 to 23. This renumbering stems from updated assembly orientation and synteny mapping, aiming to harmonize chromosome identities across species for comparative genomics (e.g. aligning acrocentric chromosomes consistently across primates). The updated representation better reflects the ancestral karyotype of great apes, improving cross-species comparisons by anchoring each chromosome to shared structural homology rather than legacy identifiers (Mohanty et al., 2025; Yoo et al., 2025).

Despite the availability of newer telomere-to-telomere (T2T) genome assemblies for *Pongo abelii* and *Pongo pygmaeus*, we have retained Susie_PABv2 and its corresponding chromosome numbering (14 for IGH, 2A for IGK, and 22 for IGL) as the reference framework for IG loci annotation in the IMGT database. Susie_PABv2 remains widely used reference for *Pongo abelii* (Makova et al., 2024; Olivieri & Gambón Deza, 2018; Zhou et al., 2023), and it is the assembly upon which prior IMGT locus definitions and gene nomenclature have been based. Maintaining consistency with this reference ensures compatibility with existing datasets and allows for direct comparisons across species and individuals without introducing ambiguity due to shifts in chromosome identifiers. However, in analyses and database integration involving the newer T2T assemblies, we adopt the updated chromosome numbers (e.g., IGH on chromosome 15, IGK on 12, IGL on 23). This dual approach preserves continuity with established immunogenetic references while also recognizing the improved structural accuracy of recent assemblies, allowing us to bridge historical annotations with evolving genomic representations.

### 3.2 Genetic structure and localization of IG loci

In *Pongo abelii*, the immunoglobulin (IG) loci are located as follows: the IGH locus is on chromosome 14 in the Susie_PABv2 assembly and on chromosome 15 in the newer NCBI genome assemblies; the IGL locus is on chromosome 22 in Susie_PABv2 and on chromosome 23 in the new assemblies; and the IGK locus is on chromosome 2A in Susie_PABv2 and on chromosome 12 in the new assemblies. In *Pongo pygmaeus*, where annotations are done only in the newer NCBI genome assemblies, the IGH, IGL, and IGK loci are located on chromosomes 15, 23, and 12, respectively. These differences do not reflect biological rearrangements but rather revisions in chromosome naming and organization following improvements in sequencing accuracy, scaffolding, and genome alignment. Essentially, the IG loci occupy the same physical regions in the genome, but their chromosomal identifiers have been renumbered to align with the updated assembly structure.

#### 3.2.1 IGH: Immunoglobulin Heavy Chain in *Pongo* Species

The IGH locus is located immediately adjacent to the telomere on the long (q) arm comprising a total of 161 IGHV genes and 274 alleles in *Pongo abelii* and 177 genes and 259 alleles in *Pongo pygmaeus* were annotated. Distribution of the genes within the IGHV subgroups was not even, similar to that observed in other species. Most of the IGHV genes belong to the IGHV3 subgroup. This subgroup is also the most prevalent in rhesus macaque and human IGH annotation (Lefranc, 2001; Nguefack Ngoune et al., 2022; Papadaki et al., 2025). The other IGHV subgroups were represented by a comparatively low number of genes. Additionally, a difference in gene content among different *Pongo* assemblies was found in Figure 1A and B. Alignment of the functional, ORF and pseudo in frame IGHV genes, is provided in (https://www.imgt.org/IMGTrepertoire/Proteins/proteinDisplays.php?species=Sumatran%20orangutan&latin=Pongo%20abelii&group=IGHV) and in (https://www.imgt.org/IMGTrepertoire/Proteins/proteinDisplays.php?species=Bornean%20orangutan&latin=Pongo%20pygmaeus&group=IGHV).

**Figure 1.**
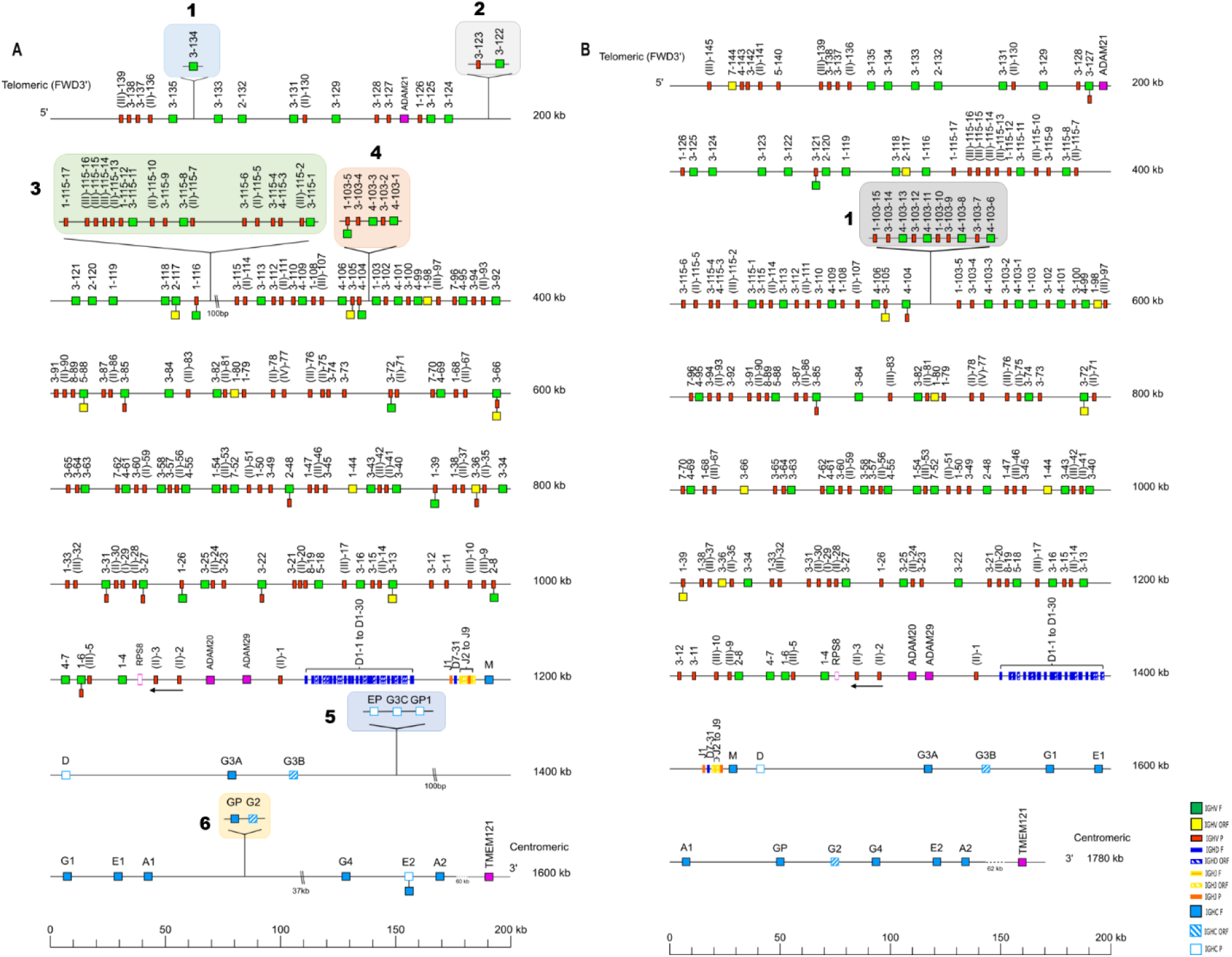
Holistic map of the immunoglobulin heavy chain (IGH) locus in (A) *Pongo abelii* and (B) *Pongo pygmaeus.* The locus organization is deduced from 3 chromosome-localized genome assemblies in *Pongo abelii* and 2 in *Pongo pygmaeus*. Genes are shown in their relative positions across the locus.The diagram shows the IGH genes and positions on the locus according to the IMGT nomenclature. The arrows indicate an inverse transcriptional orientation in the locus. Shaded regions represent haplotype-specific insertions, deletions, or structural variants, validated by comparison among haplotypes and individuals from both *Pongo* species and supported by read alignments visualized in IGV.

Nine IGHJ genes and 13 alleles in *Pongo abelii* and 9 IGHJ genes and 12 alleles were identified and localized in the *Pongo pygmaeus* IGH locus. The organization of the IGHJ genes is comparable to that of the human J-CLUSTER. They were classified into nine subsets, all named after their human counterparts (Figure 1A and B). Nucleotide and amino acid alignment of the functional, ORF and in frame pseudo genes IGHJ genes is available (https://www.imgt.org/IMGTrepertoire/Proteins/alleles/index.php?species=Pongo%20abelii&group=IGHJ&gene=IGHJ-overview) (https://www.imgt.org/IMGTrepertoire/Proteins/alleles/index.php?species=Pongo%20pygmaeus&group=IGHJ&gene=IGHJ-overview). The reading frame of the functional IGHJ genes encodes the expected motif WGXG (tryptophan, glycine, any amino acid, and glycine).

31 IGHD genes belonging to 7 sets were identified and localized in both orangutan IGH locus, comprising 35 alleles in *Pongo abelii* and 34 alleles in *Pongo pygmaeus*. Each IGHD set comprises seven genes except for the IGHD1 set which is the most represented with six genes and IGHD7 set which is the least represented with one gene Figure 1A and B. Alignment of the functional, ORF and pseudo in frame IGHD genes, is provided in https://www.imgt.org/IMGTrepertoire/Proteins/alleles/index.php?species=Pongo%20abelii&group=IGHD&gene=IGHD-overview and in https://www.imgt.org/IMGTrepertoire/Proteins/alleles/index.php?species=Pongo%20pygmaeus&group=IGHD&gene=IGHD-overview. A 20X zoom view of the D-J-CLUSTER is provided in Supplementary Figure 3.

Thirteen constant IGHC genes corresponding to 29 alleles encoding the constant region of the H chain isotypes {(H-mu, H-delta, H-gamma1 (x 3), H-gamma2, H-gamma3 (x 3), H-gamma4, H-epsilon (x3), and H-alpha(x 2)} and associated exons were mapped onto assemblies of *Pongo abelii* with IGHGP1, IGHEP and IGHG3C only found in reference assembly (GCA_002880775.3). Likewise, twelve constant IGHC genes corresponding to 17 alleles encoding the constant region of the H chain isotypes {(H-mu, H-delta, H-gamma1 (x 2), H-gamma2, H-gamma3 (x 2), H-gamma4, H-epsilon (x2), and H-alpha(x 2)} and associated exons were mapped onto assemblies of *Pongo pygmaeus* Figure 1A and B.

The TMEM121 gene is located immediately downstream of the IGH locus at its 5’ end, approximately 62 kb beyond the terminal constant gene IGHA2. This position places TMEM121 within proximity to the 3′ regulatory region (3′RR), a cluster of powerful enhancers that control class-switch recombination and antibody expression in B cells. Because of this spatial relationship, TMEM121 is considered an IGH-borne gene potentially influenced by the IGH regulatory architecture (Birshtein, 2014).

#### 3.2.2 IGL: Immunoglobulin Lambda Chain in *Pongo* Species

The IGL is organized into a cluster of variable (IGLV), joining (IGLJ), and constant (IGLC) genes. A total of 105 IGLV genes corresponding to 198 IGLV alleles in *Pongo abelii* and 114 IGLV genes corresponding to 172 IGLV alleles in *Pongo pygmaeus* were annotated. The IGLV genes were classified into 11 subgroups and seven clans defined according to IMGT-ONTOLOGY and to their sequence similarity with the human IGLV subgroups (https://www.imgt.org/IMGTindex/IGLVclans.php). The orangutans IGL J-C-CLUSTER is composed of seven tandems of IGLJ and IGLC genes.

The VPREB1 (pre-B lymphocyte 1) gene is situated within the immunoglobulin lambda IGL locus, positioned upstream of the lambda variable (IGLV) gene cluster. It encodes a small immunoglobulin-like protein that partners with λ5 (IGLL1) to form the surrogate light chain component of the pre-B-cell receptor, which plays an essential role in early B-cell differentiation and selection (Sabbattini & Dillon, 2005).

The locus is flanked by two conserved non-immunoglobulin “borne” genes. The 5′ borne gene, TOP3B (DNA topoisomerase III beta), lies immediately upstream of the most distal IGLV gene, whereas the 3′ borne gene, RSPH14 (radial spoke head 14 homolog), is located downstream of the terminal constant gene IGLC. In humans, TOP3B and RSPH14 are positioned approximately 43 kb and 136 kb from the first and last IGL genes, respectively. In orangutans these borne distances are expanded relative to humans: TOP3B is located 61 kb upstream of the first IGLV gene in *Pongo abelii* and 91 kb in *Pongo pygmaeus*, while RSPH14 resides 128 kb and 219 kb downstream of the terminal IGLC gene in the same species, respectively.

Within the IGL locus, several region-positioned intervening (RPI) genes including PRAME, ZNF280A, and ZNF280B were also identified. These are non-immunoglobulin genes integrated within the IGL locus rather than flanking it.

#### 3.2.3 IGK: Immunoglobulin Kappa Chain in *Pongo* Species

The IGK locus consists of a cluster of variable (IGKV) genes, joining (IGKJ) genes, and a single constant (IGKC) gene. IGK locus in *Pongo abelii* comprises 79 IGKV genes corresponding to 150 IGKV alleles, 5 IGKJ genes and alleles and one IGKC gene and allele. IGK locus in *Pongo pygmaeus* harbours 77 IGKV genes corresponding to 124 IGKV alleles, 5 IGKJ genes corresponding to 6 IGKJ alleles and one IGKC gene and 2 IGKC alleles.

The IGK locus is flanked by two conserved non-immunoglobulin “borne” genes: PAX8 (5′) upstream of the most distal IGKV gene and RPIA (3′) downstream of the terminal IGKC gene. In *Pongo abelii*, PAX8 is located 312 kb upstream and RPIA 136 kb downstream of the IGK gene cluster, whereas in *Pongo pygmaeus*, PAX8 lies 260 kb upstream and RPIA 132 kb downstream of the corresponding IGK region.

### 3.3 IG Loci Structure and Gene Functionality Across Five *Pongo* Assemblies

To investigate further the structural diversity and gene composition of the IGH locus across the assemblies studied, we analyzed key characteristics, such as number of genes and gene functionality Supplementary Table 2-4. Figure 4 provides an overview of the gene composition within complete IGH loci across diverse *Pongo* assemblies. The gene composition and functionality classification of the complete immunoglobulin loci—IGK (A), IGL (B), and IGH (C)—were examined across five chromosome-localized *Pongo* assemblies. All three loci were presented on the same genomic scale to allow direct comparison of locus size, gene number, and functional composition.

Each locus was analyzed according to IMGT gene functionality categories, including functional, open reading frame (ORF), and pseudogene classes. The stacked horizontal bars illustrate the proportional distribution of each functionality type per assembly and per locus, while the accompanying pie charts summarize the functionality proportion of variable genes across assemblies.

Across all loci, pseudogenes formed the largest proportion of variable genes, particularly in the IGH locus (Figure 2 C), which exhibited a higher number of pseudogenes and overall locus expansion compared to the light chain loci (IGK and IGL). Functional genes and ORFs comprised smaller fractions, with relative proportions varying between loci. Specifically, the IGH locus contained approximately 52–70% pseudogenes, whereas the IGK and IGL loci displayed higher relative proportions of functional genes (28–42%) and only minor contributions from ORFs (2–6%). The J, D, and C regions are comparatively conserved, showing little or no variation across different genomes, suggesting strong selective maintenance of their structural and functional integrity.

**Figure 2.**
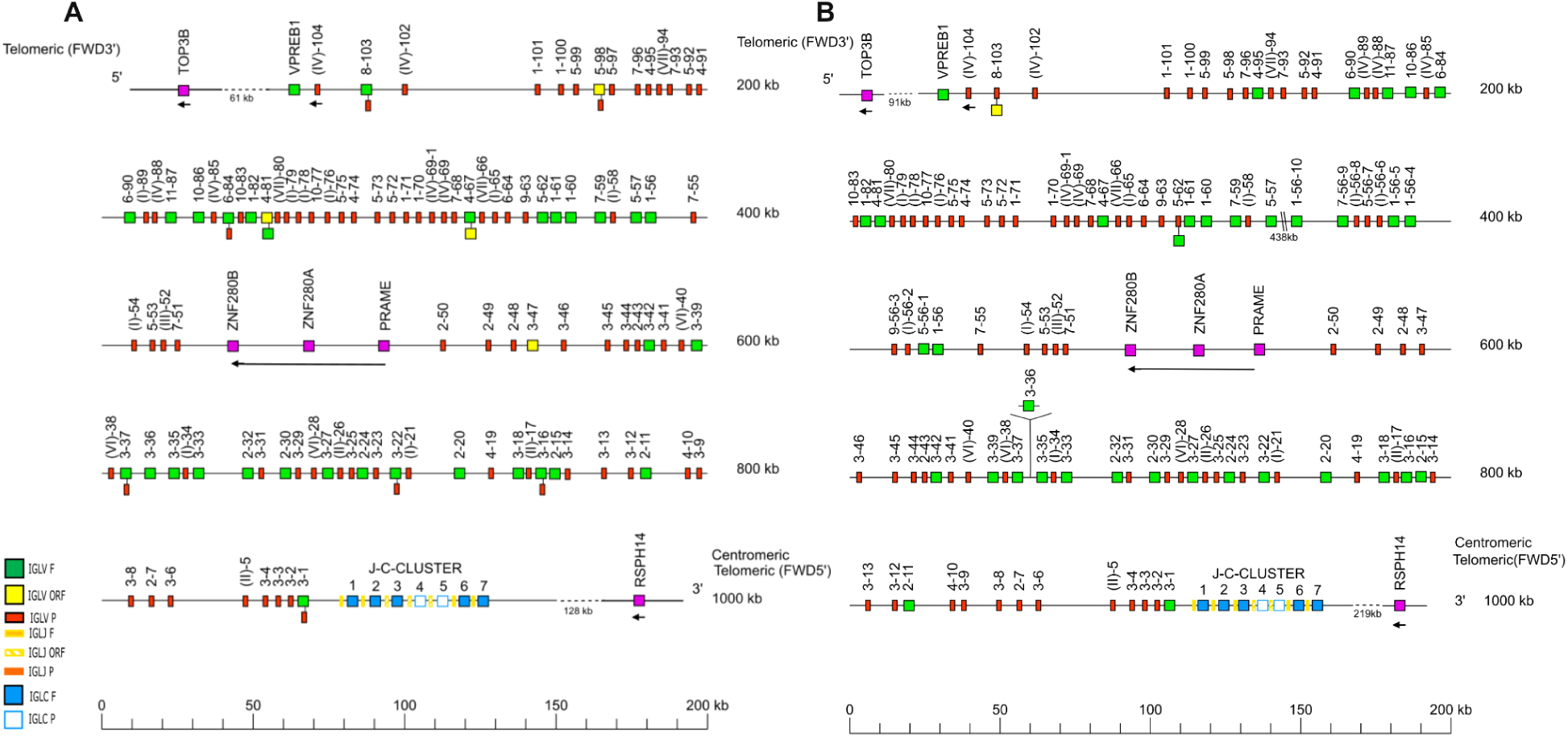
Holistic map of the immunoglobulin Lambda light chain (IGL) locus in (A) *Pongo abelii* and (B) *Pongo pygmaeus.* The locus organization is deduced from 3 chromosome-localized genome assemblies in *Pongo abelii* and 2 in *Pongo pygmaeus*. Genes are shown in their relative positions across the locus.The diagram shows the IGH genes and positions on the locus according to the IMGT nomenclature. The arrows indicate an inverse transcriptional orientation in the locus. The map highlights differences in gene functionality among alleles of the same IGL genes.

The consistent scale across panels highlights clear differences in locus size, with the IGH locus being substantially larger and functionally more diverse, indicating an evolutionary expansion of the heavy chain repertoire relative to the more compact light chain loci.

### 3.4 Structural variations/haplotype-specific insertions and duplications

The high frequency of structural variation observed across the IGHV locus is thought to be driven by its repetitive nature, largely caused by the duplication of IGHV genes (Pramanik et al., 2011). To explore this structural complexity in detail, holistic maps of the immunoglobulin heavy chain (IGH) locus in *Pongo abelii* and *Pongo pygmaeus* were constructed from localized genome assemblies to illustrate the overall organization and structural variation within the locus (Figure 1A and B). The locus shows distinct shaded regions Figure A(1–4) and B (1), each representing a structurally or functionally unique segment characterized by haplotype-specific insertions or deletions. The boundaries and composition of each region were validated through comparison across multiple haplotypes, assemblies, and individuals from both *Pongo abelii* and *Pongo pygmaeus*, and were further supported by ultra long read alignments visualized in IGV (Supplementary Figures 5-9).

The telomeric end of the IGH locus in *Pongo pygmaeus* (Figure 1B) shows a high degree of conservation with the human and gorilla IGHV gene cluster. In *Homo sapiens, Gorilla gorilla gorilla* and *Pongo pygmaeus*, this region contains a series of IGHV genes belonging to the same IGHV subgroups, exhibiting similar functionality. In the human and gorilla, this telomeric cluster includes genes such as IGHV5-78, IGHV(II)-78-1, IGHV3-79, IGHV4-80, IGHV7-81, and IGHV(III)-82, which are also represented in *Pongo pygmaeus* with conserved order and orientation. The gene content and order are remarkably conserved, indicating a shared ancestral organization of the telomeric IGHV domain among great apes. In *Pongo abelii*, however, the locus terminates at IGHV(III)-139, representing a slightly shorter configuration (Figure 5).

Regions 1 and 2 include the genes IGHV3-134 and IGHV3-123/IGHV3-122, which are present in some *Pongo* assemblies but absent in others Supplementary Figure 4. This variable presence suggests a small-scale structural polymorphism within this region.

Region 3 contains a dense cluster of IGHV genes between genes IGHV3-115 and IGHV1-119. The number of genes in this region differ among the *Pongo abelii* assemblies, showing clear variation in gene content.

Region 4 is located between IGHV1-98 and IGHV4-104 and consists of tandem repeats of a five-gene block. The first block spans IGHV4-99 through IGHV1-103, and this same five-gene block is duplicated multiple times. These duplications give rise to IGHV4-103-1 through IGHV4-103-15, representing three consecutive repetitions of the same five-gene structure, each retaining the same IGHV subgroup composition. The number of repeated blocks is not constant and varies across assemblies. In the NHGRI_mPonPyg2-v2.0_pri (primary haplotype) (Figure 6B), only a single instance of this block is present, spanning IGHV4-103-1 through IGHV4-103-5. In contrast, the NHGRI_mPonPyg2-v2.0_alt (alternate haplotype) (Figure 6A) shows a much larger expansion, extending continuously to IGHV4-103-15, corresponding to three full consecutive copies of the same five-gene structure.

**Figure 3.**
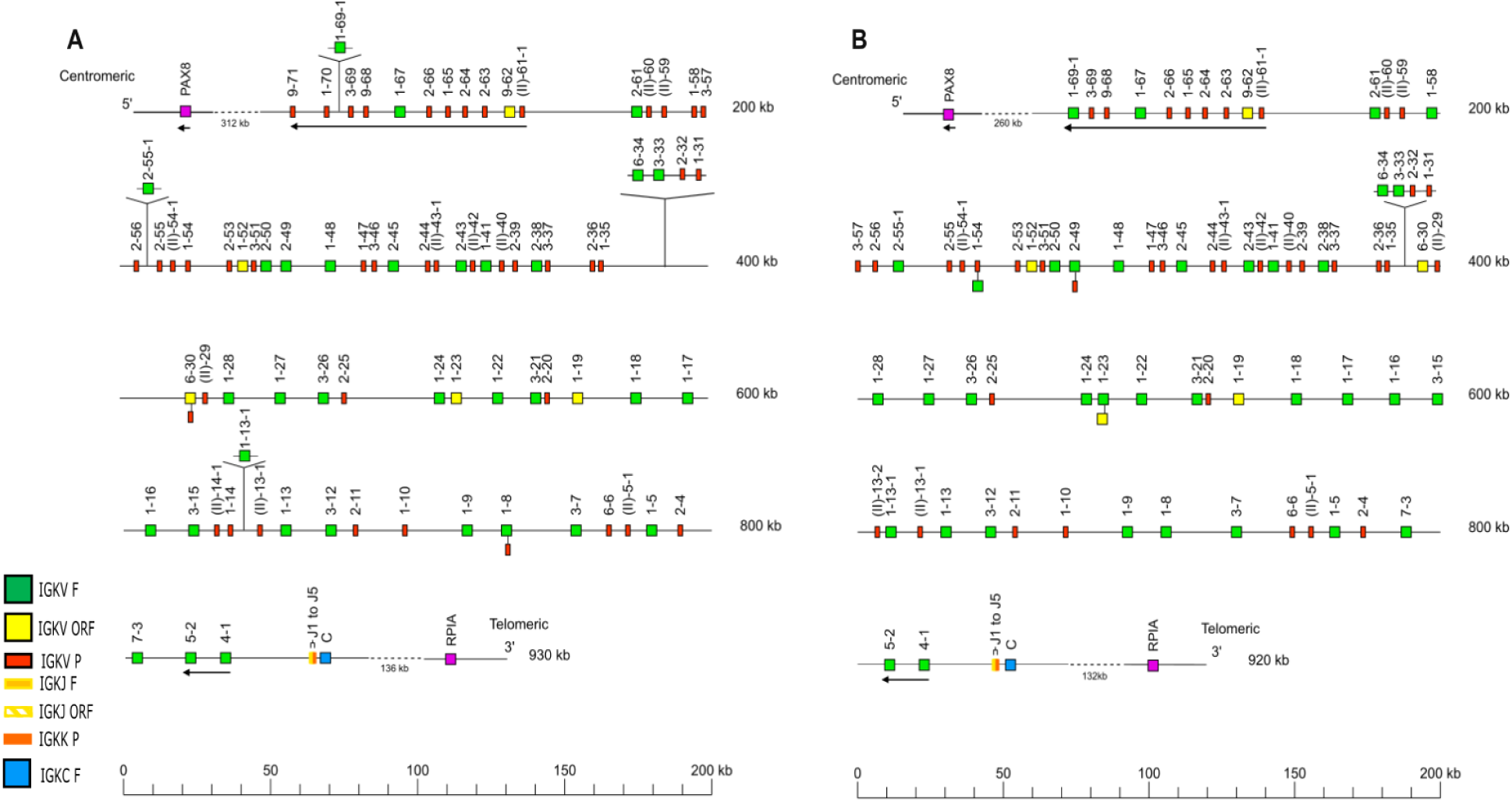
Holistic map of the immunoglobulin Kappa light chain (IGK) locus in (A) *Pongo abelii* and (B) *Pongo pygmaeus.* The locus organization is deduced from 3 chromosome-localized genome assemblies in *Pongo abelii* and 2 in *Pongo pygmaeus*. Genes are shown in their relative positions across the locus.The diagram shows the IGK genes and positions on the locus according to the IMGT nomenclature. The arrows indicate an inverse transcriptional orientation in the locus. The map also highlights differences in gene functionality among alleles of the same IGL genes.

**Figure 4.**
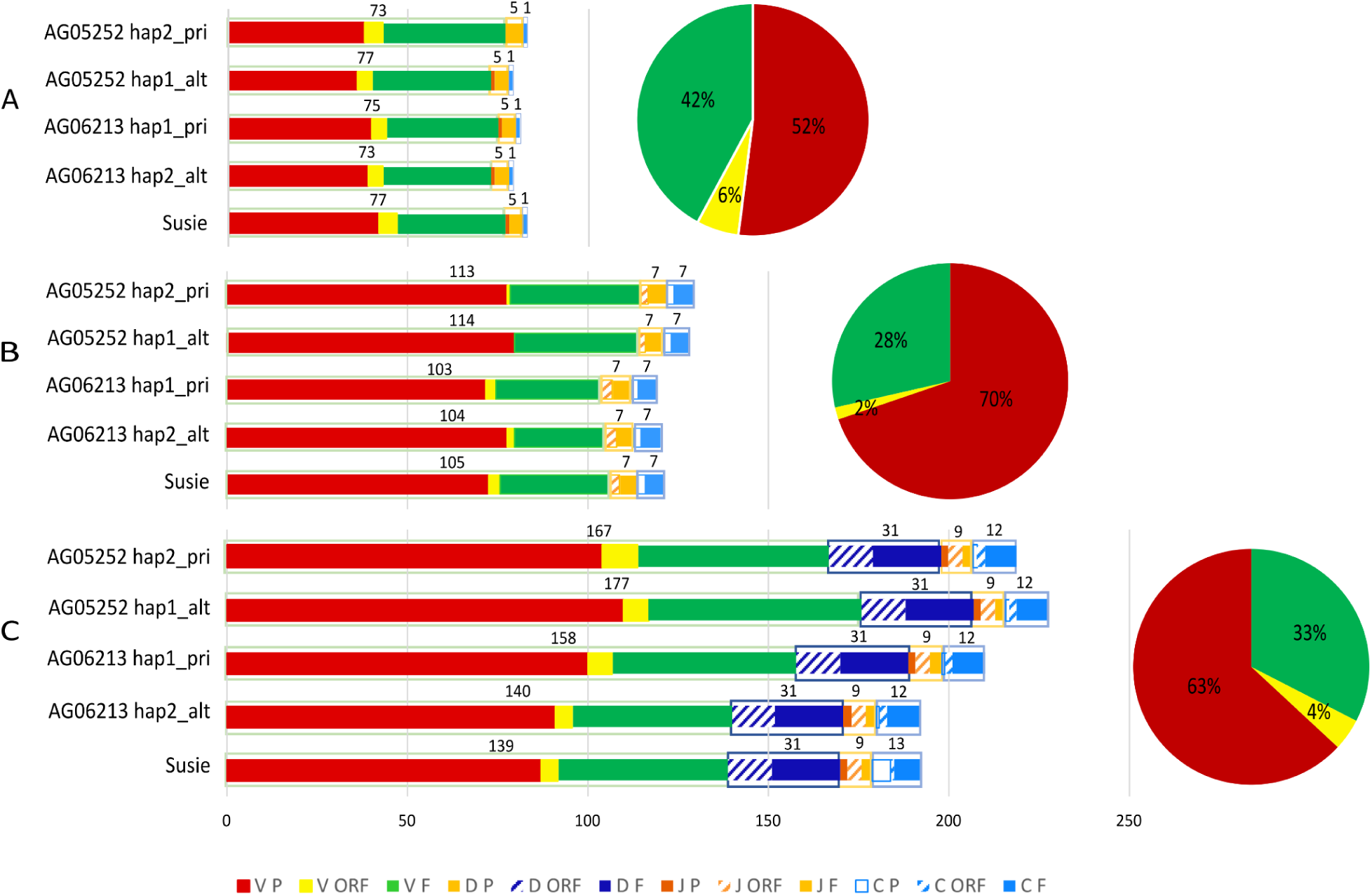
Gene composition and functionality classification of complete IG Loci IGK (A), IGL (B), and IGH (C) within 5 chromosome-localized *Pongo* assemblies. This graph displays the gene count and functionality distribution for each gene type (variable, diversity, joining and constant), with colors applied according to the IMGT color menu, reflecting the IMGT functionality per gene type. All loci are plotted on the same genomic scale, enabling direct comparison of locus length, gene number, and functional diversity. Pie charts on the right display the overall functional proportion of variable genes for each locus, revealing that pseudogenes dominate across all three loci, particularly within the IGH locus, which also exhibits the greatest locus size and gene complexity.

**Figure 5.**
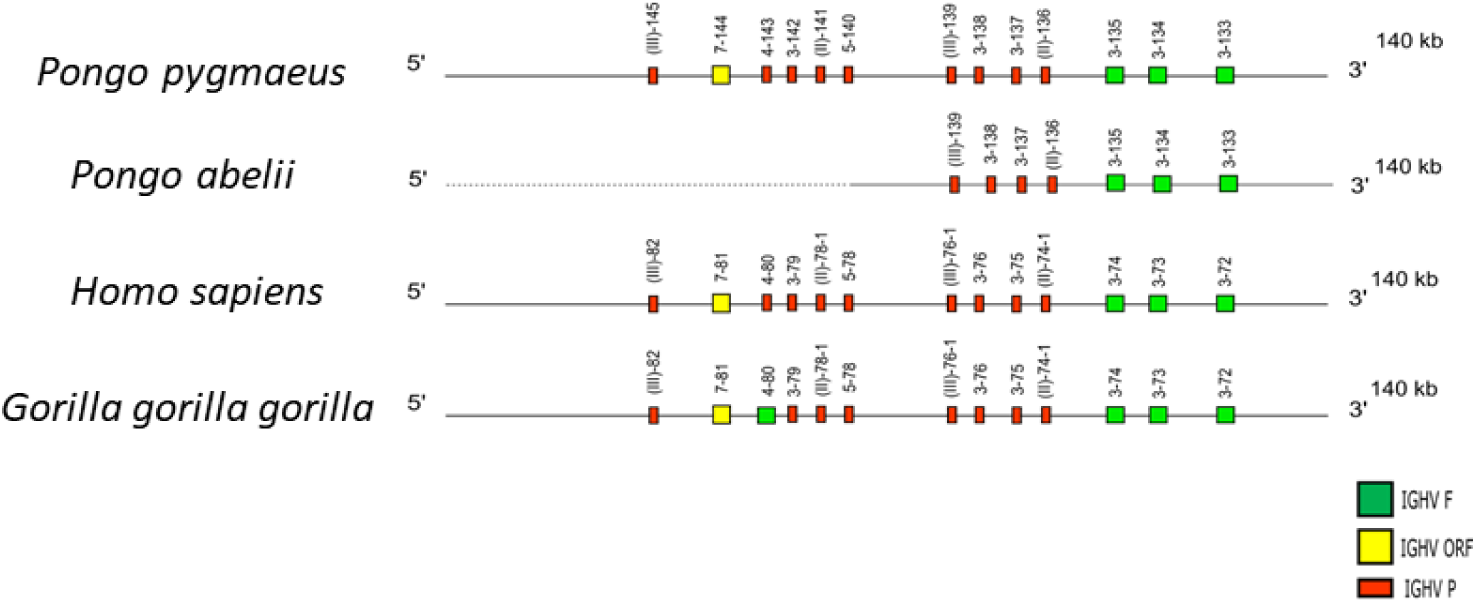
Comparative organization of the telomeric IGHV gene cluster among great apes. Schematic representation of the telomeric end of the IGH locus in *Pongo pygmaeus, Pongo abelii, Homo sapiens,* and *Gorilla gorilla gorilla*. The cluster displays a highly conserved architecture in *Pongo pygmaeus, Homo sapiens,* and *Gorilla gorilla,* reflecting strong evolutionary conservation of this telomeric region. In contrast, *Pongo abelii* displays a slightly shorter telomeric configuration.

**Figure 6.**
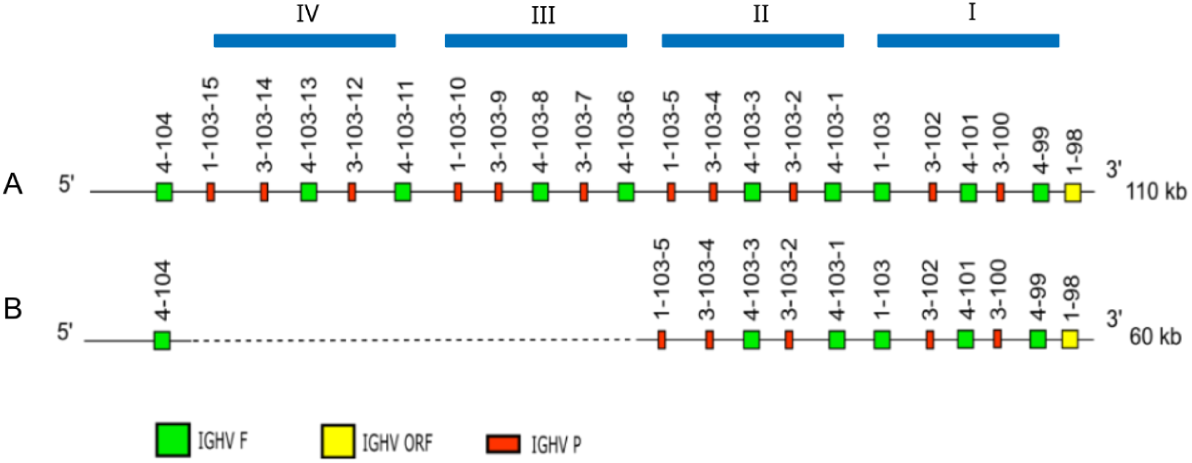
Haplotype-specific expansion of a duplicated IGHV block in Region 4. Region 4, located between IGHV1-98 and IGHV4-104, contains tandem repeats of a five-gene block that runs from IGHV4-99 through IGHV1-103. This same block is duplicated in series, giving rise to IGHV4-103-1 through IGHV4-103-15, corresponding to up to three consecutive copies with identical IGHV subgroup composition. The number of block copies varies across *Pongo* assemblies.

In the Susie_PABv2 reference assembly, this region was not fully represented, most likely due to its highly repetitive structure, which makes it difficult to resolve completely using conventional assembly methods. Analysis of the new Susie haplotype-resolved assemblies (Susie_PAB_hifiasm-v0.15.2.pri and Susie_PAB_hifiasm-v0.15.2.alt) provided additional resolution of this area: the primary haplotype contains genes IGHV4-103-1 to IGHV4-103-5, whereas the alternate haplotype extends up to IGHV4-103-10. These newer assemblies therefore complement the Susie_PABv2 reference by filling previously unresolved repetitive regions.

Regions 5 and 6 encompass the constant region (IGHC) genes of the IGH locus. In most *Pongo* assemblies, this region contains the expected IGHC repertoire, including IGHGP, IGHG2, IGHG3, IGHE, and IGHA, arranged in the canonical order and orientation. However, in the Susie_PABv2 assembly, three additional IGHC genes IGHGP1, IGHEP, and IGHG3C were annotated, while IGHGP and IGHG2 were absent.

Read mapping across this region (Supplementary Figure 10) showed clear coverage dropouts and low mapping quality precisely at the positions corresponding to IGHGP and IGHG2, suggesting these absences result from unresolved N-gap, reflecting incomplete sequence representation rather than true gene loss. Conversely, the regions annotated as IGHGP1, IGHEP, and IGHG3C coincide with irregular coverage patterns, multiple secondary alignments, and local sequence gaps, indicative of false duplications or misassembled fragments in low-complexity constant-region sequences.

Further confirmation came from the recently generated Susie_PAB_hifiasm-v0.15.2.pri and Susie_PAB_hifiasm-v0.15.2.alt assemblies of the same individual, which recovered IGHGP and IGHG2 but did not contain IGHGP1, IGHEP, or IGHG3C. Although these assemblies are not included in public databases due to not meeting IMGT quality criteria (Supplementary Table 1), their concordance with other *Pongo abelii* and *Pongo pygmaeus* individuals confirms that the three additional IGHC genes in Susie_PABv2 are assembly artifacts. Accordingly, IGHGP1, IGHEP, and IGHG3C will be flagged as false gene predictions in our dataset. Taken together, Regions 5–6 show that the constant region is generally conserved across orangutan genomes, and the apparent differences in Susie_PABv2 reflect assembly-specific anomalies rather than genuine structural variation.

### 3.5 Haplotype-Resolved Heterozygosity of Immunoglobulin Variable Gene Families in *Pongo pygmaeus* and *Pongo abelii*

To further characterize structural variation within the IG loci, we examined genes present exclusively on one haplotype, distinguishing between hemizygous deletions and haplotype-specific insertions or duplications. Hemizygous genes, defined as those present on one haplotype but completely absent on the homologous chromosome, typically reflect deletion events and result in reduced copy number. In contrast, we also identified instances where additional gene copies were found on only one haplotype, with no corresponding ortholog on the other. These haplotype-specific insertions or duplications increase gene copy number asymmetrically.

Across all three immunoglobulin loci, the distribution of homozygous, heterozygous, and hemizygous genes revealed pronounced haplotype-level diversity. The majority of genes displayed allelic heterozygosity, showing that sequence variation between shared genes is more common than gene gain or loss. Notably, the IGH locus contained the highest proportion of hemizygous genes in both *Pongo pygmaeus* and *Pongo abelii*, suggesting that this region undergoes frequent structural variation, including haplotype-specific deletions and insertions. In contrast, the IGLV and IGKV loci were dominated by heterozygous but largely balanced allelic variants, consistent with stronger sequence conservation and lower rates of copy-number turnover. Comparative analysis between the two species further showed that *P. abelii* exhibits slightly higher overall gene-level heterozygosity across all three loci, indicating greater allelic diversification within its immunoglobulin repertoire. Together, these results highlight extensive allelic and structural heterogeneity within orangutan IG loci, with the IGH locus emerging as a hotspot for haplotype-specific structural variation.

### 3.6 Switch sequence analysis in IGHC genes

To investigate the regulatory architecture of orangutan immunoglobulin heavy chain constant (IGHC) genes, we first extracted the 5 kb strand-aware upstream switch-region sequences for each constant gene from the *Pongo pygmaeus* and *Pongo abelii* reference genomes (GCF_028885625.2 and GCF_028885655.2, respectively). We quantified the density of canonical AID hotspot motifs including GAGCW (W = A or T) and GGGGBT (B = C, G, or T) and calculated the GC content across the same regions (Supplementary Table 5). When visualized as bar charts, these features showed a highly concordant and gene-specific enrichment pattern between the two species, indicating strong conservation of switch-region architecture across orangutans (Figure 8). Quality control (QC) plots showing sliding-window distributions of motif density and GC fraction were generated for each region to verify extraction accuracy and to visualize local enrichment patterns in both *Pongo* species Supplementary Figure 12. To further explore the structural organization underlying this conservation, we generated Gepard self-alignment dotplots for *Pongo abelii*, both across the entire IGHC constant-gene cluster and individually for each upstream switch-region (Krumsiek et al., 2007).

**Figure 7.**
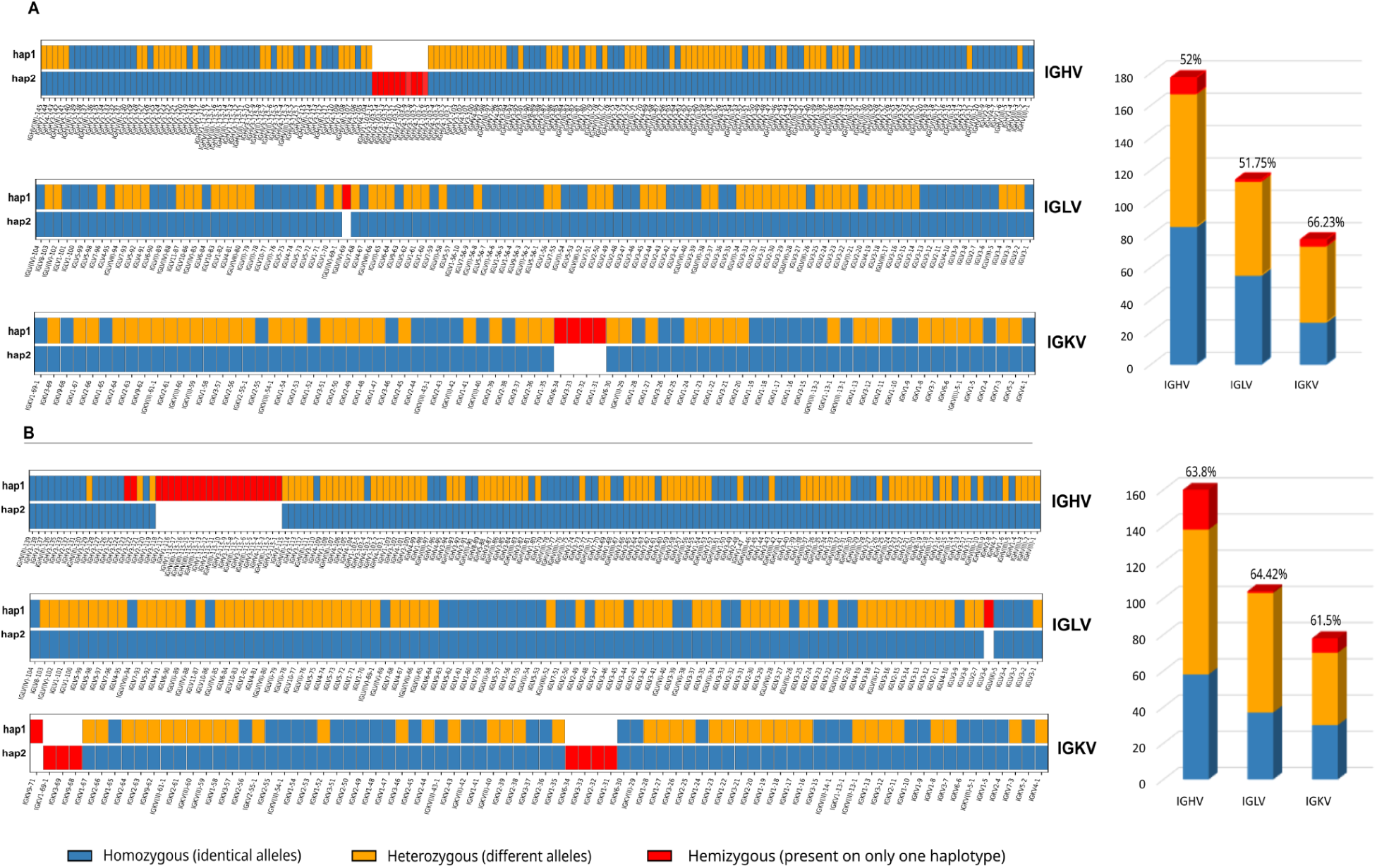
Haplotype-resolved heterozygosity across immunoglobulin variable loci in orangutans. Panels (A) and (B) show comparisons of *Pongo pygmaeus* and *Pongo abelii* haplotypes, respectively, for the immunoglobulin heavy (IGHV), lambda (IGLV), and kappa (IGKV) variable gene families. Each horizontal track represents one haplotype (hap1 and hap2) aligned gene-by-gene. The colors indicate the degree of allelic divergence at each gene locus: blue denotes homozygous genes that are identical in both haplotypes, orange denotes heterozygous genes present on both haplotypes but differing in sequence (allelic variants), and red denotes hemizygous genes present on only one haplotype. Bar charts to the right of each panel summarize the proportions of the three categories for each locus, with percentage values representing overall gene-level heterozygosity. The figure illustrates that both orangutan species exhibit substantial allelic and structural variation across the IGHV, IGLV, and IGKV loci, with *P. abelii* showing slightly higher overall heterozygosity than *P. pygmaeus.* Assemblies used: NHGRI_mPonPyg2-v2.- (pri/alt) and NHGRI_mPonAbe1-v2.- (pri/alt).

**Figure 8:**
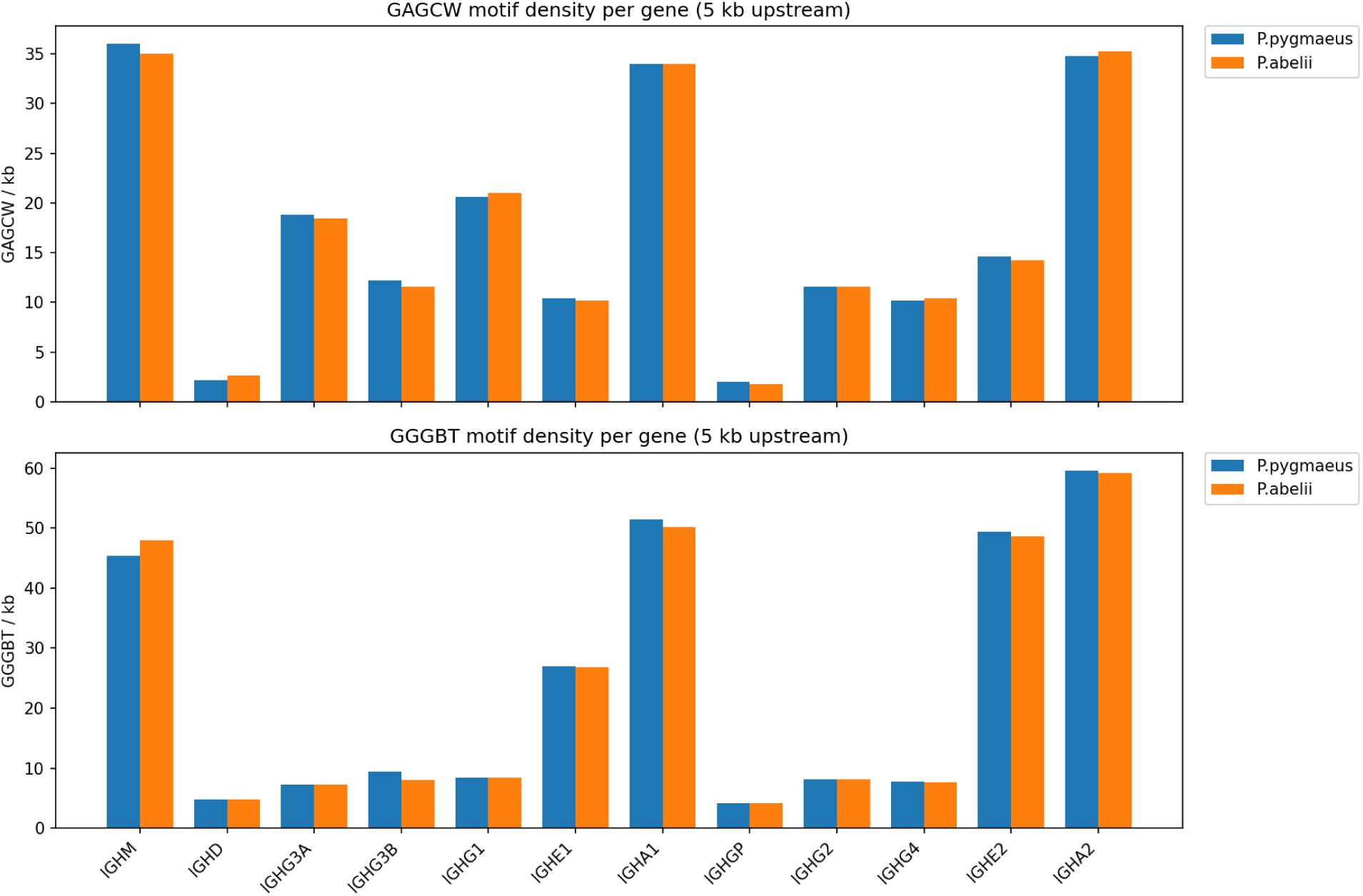
Density of canonical AID hotspot motifs in the 5 kb upstream promoter region of each IGH constant-region gene in *Pongo pygmaeus* and *Pongo abelii*. Each bar represents the number of predicted AID target motifs per kilobase immediately upstream of the indicated IGH constant gene. The top panel shows the density of GAGCW motifs, and the bottom panel shows GGGGBT motifs. Blue bars correspond to *P. pygmaeus* and orange bars to *P. abelii*. Although absolute densities vary across constant-region subclasses, both species show a broadly conserved ranking with notably high motif enrichment upstream of IGHA1, IGHE, and IGHG2/IGHG3, and minimal enrichment at IGHD and IGHG3/IGHG4.

Distinct patterns were observed among genes corresponding to their known switch-region strength Supplementary Figure 13. The IGHM, IGHA1, and IGHA2 upstream regions displayed dense, continuous diagonals indicative of extensive tandem repeats, consistent with the canonical Sμ and Sα regions. In contrast, the IGHG and IGHE genes showed shorter or fragmented repeat tracts, reflecting more heterogeneous and less extensive switch regions. The IGHD and IGHGP upstream regions lacked internal homology signals beyond the self-identity diagonal, suggesting the absence of switch-like repeats. Overall, these per-gene dotplots confirmed the patterns seen at the cluster level Figure 9, demonstrating strong repeat enrichment upstream of μ and α genes, intermediate organization in γ and ε, and no discernible switch structures in δ or GP.

**Figure 9:**
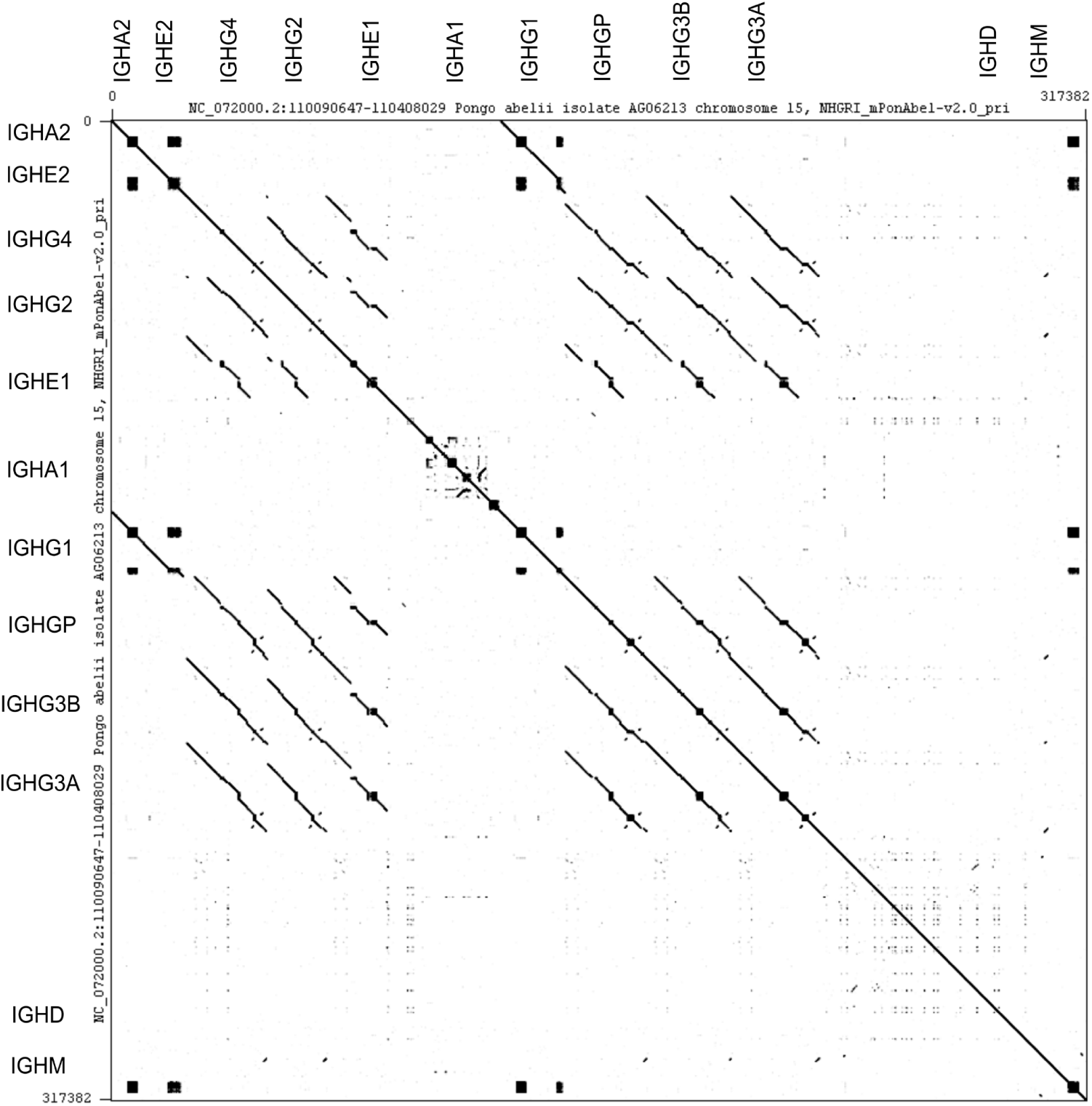
Dotplot of the IGHC locus in *Pongo abelii*. Self-comparison of the constant region cluster generated with standalone Gepard application (Krumsiek et al., 2007) shows extended tandem repeats upstream of IGHM and IGHA genes, consistent with classical switch regions. IGHE and IGHG genes display shorter or fragmented repeat signals, while IGHD and IGHGP lack detectable repeats. We used a word size of 14 because it is sufficiently small to detect the short, high-identity repeats characteristic of immunoglobulin switch regions, while still suppressing background noise from low-complexity sequences.

### 3.7 Analysis of Recombination signal sequences

Sequence logo analysis of recombination signal sequences (RSSs) from functional IG V, D, and J genes revealed the expected conserved motifs characteristic of RAG-mediated recombination Figure 10. Across all three loci (IGH, IGK, and IGL), the canonical heptamer (CACAGTG) and nonamer (ACAAAAACC) motifs were strongly conserved, consistent with previous reports (Gellert, 1997b; Hassanin et al., 2000; Nguefack Ngoune et al., 2022; Ott et al., 2025; Sirupurapu et al., 2022). The heptamer showed nearly invariant conservation at positions 1–7, particularly the central CACAGTG core, while the nonamer displayed high A content. The first three bases (CAC) of the heptamer and the central A-rich core of the nonamer are critical for RAG1/2 recognition and cleavage during V(D)J recombination. Minor locus-specific variations were observed at the flanking nucleotides and within the spacer regions, suggesting subtle differences in recombination efficiency or RAG binding preferences. The spacer length distribution confirmed the expected predominance of 12 bp and 23 bp spacers, validating the integrity of the extracted sequences. Together, these results confirm that the RSS motifs across functional IG genes retain the canonical sequence features necessary for efficient V(D)J recombination. Figure 10 shows the consensus RSS motifs in *Pongo abelii*, while Supplementary Figure 11 presents the corresponding motifs for *Pongo pygmaeus*.

**Figure 10.**
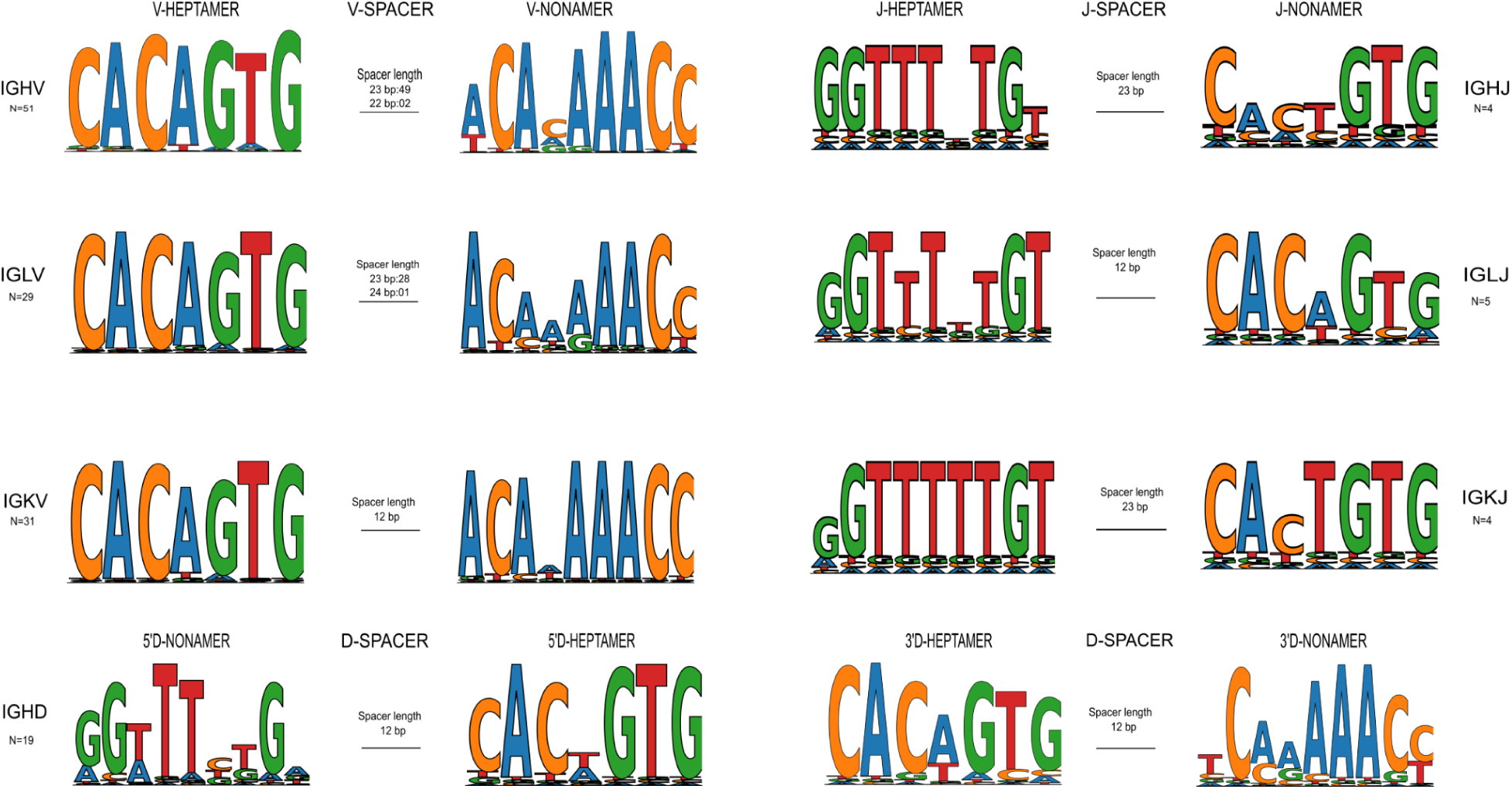
Consensus motifs of recombination signal sequences (RSSs) from functional immunoglobulin genes in *Pongo abelii*. Sequence logos derived from functional IG V, D, and J genes highlight the highly conserved heptamer (CACAGTG) and nonamer (ACAAAAACC) motifs, which flank coding segments and direct RAG-mediated recombination. Both motifs show strong positional conservation, particularly within the core CAC and ACA elements, consistent with their essential role in synapsis and cleavage. The observed spacer length distribution confirmed the expected 12/23-bp rule, with minor locus-specific sequence variability suggesting potential fine-tuning of recombination efficiency across loci.

### 3.8 Phylogeny of IG V genes

Across genome assemblies of *Homo sapiens, Pongo abelii, Pongo pygmaeus* and *Gorilla gorilla gorilla*, we quantified the immunoglobulin heavy-chain (IGHV) and light-chain (IGKV, IGLV) V-CLUSTERS by taking into account the total V-CLUSTER length and the number of annotated V genes (Supplementary Table 6).

The mean gene density values revealed clear locus- and species-specific patterns. Among all four primates, the IGHV locus showed the greatest divergence, with *Homo sapiens* exhibiting the highest density, indicating a more compact and gene-rich heavy-chain V-CLUSTER relative to the other species. In contrast, *Gorilla gorilla gorilla* displayed a noticeably lower IGHV density, reflecting a more expanded but less gene-dense organization of this locus Figure 11 A.

**Figure 11.**
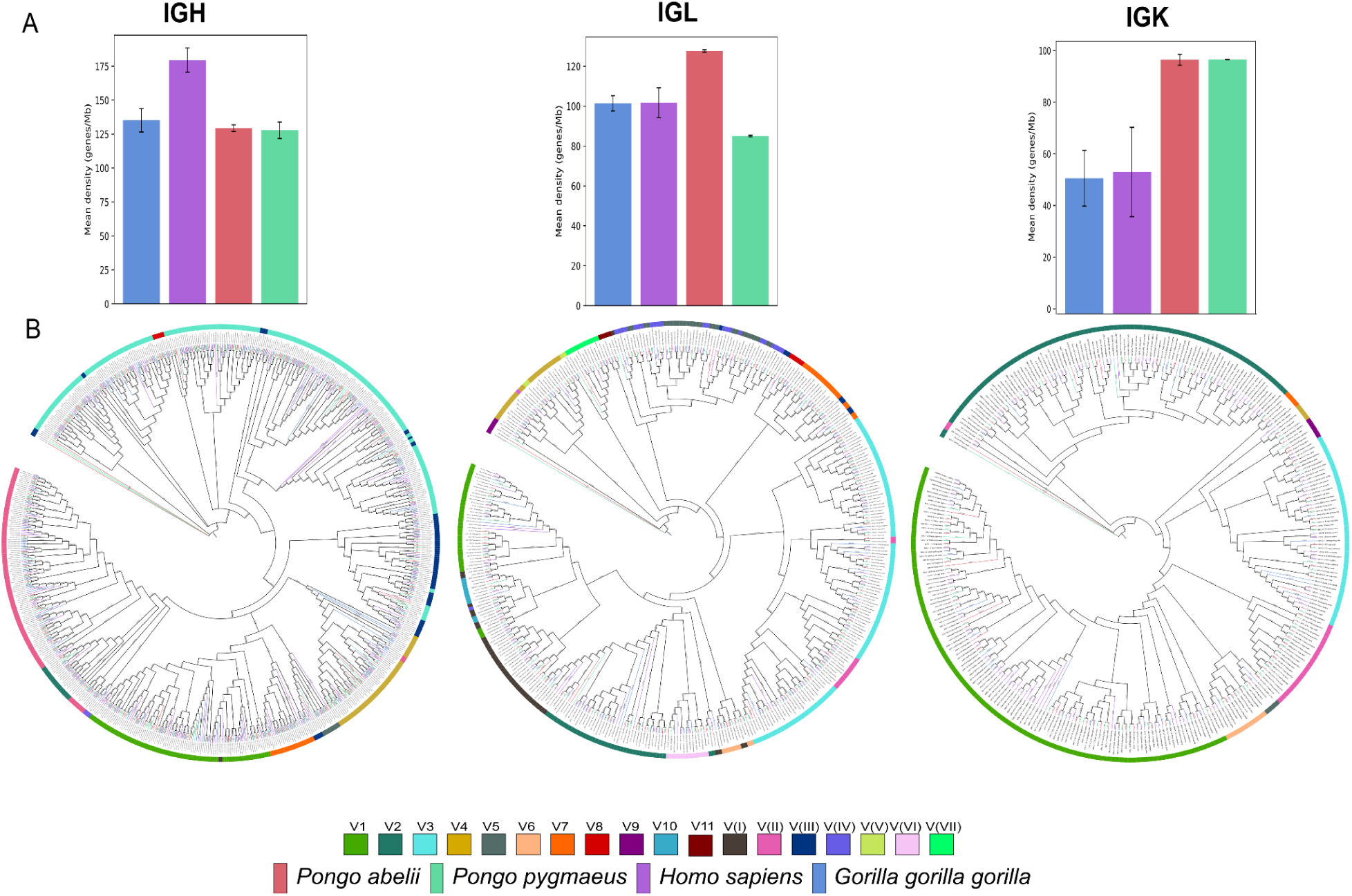
Comparative analysis and phylogenetic relationships of immunoglobulin variable (V) genes across three loci in great apes. (A) Gene density distributions (genes/Mb) for the IGHV, IGKV, and IGLV loci in *Homo sapiens, Pongo abelii, Pongo pygmaeus*, and *Gorilla gorilla gorilla*. Each bar represents the mean gene density ± SD across genome assemblies, showing interspecies variation in V-gene organization. (B) Maximum likelihood phylogenies of IGHV, IGKV, and IGLV V genes, color-coded by species and subgroup. Phylogenetic clustering reveals conserved subgroup structures with lineage-specific expansions and species-specific divergence patterns across loci.

The IGKV locus consistently showed the lowest density across all species, suggesting that this light-chain locus is generally more dispersed. However, *Pongo abelii* showed the highest density within IGKV, indicating a relatively compact organization specific to this orangutan lineage Figure 11 A.

For the IGLV locus, *Pongo abelii* again demonstrated the highest density, in contrast to *Homo sapiens*, which showed a more moderate density, suggesting that orangutans may favor a more compact light-chain organization, whereas humans prioritize a denser heavy-chain locus instead Figure 11 A.

These trends indicate that locus compaction is not uniform across primate species and that heavy- and light-chain loci follow distinct evolutionary trajectories, with *Homo sapiens* optimizing IGHV density, while orangutans favor higher density in light-chain loci, particularly IGLV.

To explore the evolutionary relationships of immunoglobulin genes across primates, we conducted a comparative analysis focusing on the 01 alleles of IGHV genes shared among *Pongo abelii*, *Pongo pygmaeus*, *Homo sapiens*, and *Gorilla gorilla gorilla*. Multiple sequence alignments and phylogenetic tree reconstructions were performed to assess sequence divergence and infer evolutionary relationships within and across these subgroups. The resulting phylogeny revealed clear clustering by IGV subgroups, with orthologous genes from different species grouping closely together, reflecting a high degree of evolutionary conservation. Phylogenetic reconstruction of all annotated V genes Figure 11 B showed conserved subgroup architectures across species, with clear clustering of orthologous genes and distinct species-specific clades. The IMGT clan reference sequences are consistently clustered within their respective subgroups, confirming accurate phylogenetic placement and evolutionary correspondence across loci. IMGT clans represent ancient evolutionary lineages of immunoglobulin variable (V) genes that share conserved framework motifs and structural features. Their inclusion provides an evolutionary reference framework, ensuring accurate subgroup classification and enabling consistent cross-species comparisons of V-gene diversification.

## 4. Discussion

Advances in large-scale DNA sequencing have transformed the study of nonhuman primate genomes. Although much genomic research has centered on human disease, there is growing interest in comparative primate genomics to both illuminate human biology and reconstruct evolutionary history. High-quality genome assemblies from great apes, including chimpanzees, bonobos, gorillas and orangutans, have already revealed unexpected insights into speciation, divergence and the mechanisms shaping genome architecture. As genomic resources continue to expand, our understanding of primate genome content, diversity and evolution is rapidly being refined. Within immunogenetics, this comparative framework is especially powerful, allowing lineage specific differences in gene content and structure to be resolved at highly complex loci, such as the immunoglobulin regions. Due to the challenges associated with obtaining high-quality morphological and genomic samples from critically endangered great apes, comprehensive analyses across all *Pongo* species remain limited. In this study, five genome assemblies representing two *Pongo abelii* individuals and two assemblies corresponding to a single *Pongo pygmaeus* individual were included. Currently, no genome assembly is available for *P. tapanuliensis*, which restricts direct comparisons with this recently described species. As additional genomic data become accessible, particularly for *P. tapanuliensis*, a more complete understanding of species-specific diversity and interspecies differences within the *Pongo* genus is expected to develop.

The IG loci show exceptionally rapid evolutionary turnover and appear to be less constrained than most other genomic regions, exhibiting frequent recombination, duplication, deletion, gene conversion, point mutation, and translocation (Das et al., 2012). Our mapping of the orangutan IG loci confirms that duplications and deletions are widespread across the variable gene region. Segmental duplications, which contribute to genomic variation in humans, also play a crucial role in driving evolutionary change across primate genomes. Approximately 5% of the human and chimpanzee genomes, and around 3.8% of the orangutan genome, are composed of these duplicated segments (Locke et al., 2011; Marques-Bonet, Ryder, et al., 2009). These genomes, particularly those of humans and other great apes, are notably enriched with dispersed duplications. This enrichment is believed to stem from a period of increased duplication activity following their evolutionary split from Old World monkeys (Jiang et al., 2007; Marques-Bonet, Kidd, et al., 2009). Segmental duplications have contributed significantly to the expansion of various protein-coding gene families, often through repeated duplication events of specific sequences (Dumas et al., 2007; Gazave et al., 2011; Marques-Bonet, Ryder, et al., 2009).

Within species, significant haplotype diversity has been observed across individuals, characterized by elevated numbers of single nucleotide variants (SNVs) in both coding and noncoding regions, as well as large-scale deletions and insertions. Some of these structural variants extend over hundreds of kilobases and encompass tens of IG genes within a given haplotype (Gidoni et al., 2019; Kaduk et al., 2022; Kidd et al., 2012; Mikocziova et al., 2020; Watson et al., 2013). The pattern we observe in repeating modules of IGHV genes (e.g. the five-gene blocks in Region 4), haplotype-specific gains and losses, and allelic variation in gene functional status is concordant with expectations from models of duplication-mediated structural evolution (Gidoni et al., 2019; Kaduk et al., 2022; Kidd et al., 2012; Mikocziova et al., 2020; Watson et al., 2013). The organization of Region 4, where a five-gene IGHV block containing two functional and three pseudogene copies is duplicated in tandem, closely resembles the structural pattern reported by (Das et al., 2008), who described a two-functional and three-nonfunctional IGHV block duplication in the human lineage. Because *Pongo* diverged earlier than *Homo* and is therefore evolutionarily more ancient, our findings indicate that this duplication architecture is not specific to humans but reflects a deeper ancestral mechanism that has been retained in great apes. The observation that *Pongo* haplotypes still show active copy number variation, with one, two, or even three repeated copies of the same five-gene block, demonstrates that this is not a fixed historical event. Instead, it represents an ongoing evolutionary process that continues to shape the IGHV locus.

The telomeric difference documented in this study may reflect the proximity of the IGH locus to the telomere, as similar telomeric positioning has been reported in human, gorilla, and orangutans (Debbagh et al., 2024; Papadaki et al., 2025). Telomeric and subtelomeric regions are known to undergo frequent segmental duplication and copy-number variation (Mefford & Trask, 2002), which aligns with the shortened configuration observed in *Pongo abelii*. In contrast to *Pongo pygmaeus*, human, and gorilla, all of which retain distal IGHV genes, *P. abelii* terminates earlier and lacks these downstream IGHV members.

The locus encoding the constant regions of immunoglobulins has undergone substantial evolutionary remodeling across vertebrate lineages. These constant regions are responsible for mediating effector functions that are distinct among immunoglobulin classes. Thus, understanding the evolutionary dynamics including the expansion or loss of immunoglobulin classes is of considerable interest in the field of comparative immunology. In mammals, five immunoglobulin classes are typically conserved. With the exception of IGHD, nearly all species maintain at least one functional gene for each class. The fundamental gene structure of immunoglobulins has remained remarkably stable since their origin over 250 million years ago (Rogers et al., 2006). However, IGHD has *been lost in* certain mammalian lineages, such as the opossum, where it has been replaced by short repeat elements derived from endogenous retroviruses (ERV) and LINE1 sequences (Wang et al., 2009). In the present study, we found that the IGHD gene is pseudogenized across all examined orangutan genomes due to the frameshifts in the exonic region. This finding is particularly notable given the phylogenetic proximity of orangutans to humans. In humans, IgD is an antibody present at very low concentrations in serum; nevertheless, it is widely expressed on the surface of B cells as part of the B cell receptor complex in many species (Dirks et al., 2023; Gutzeit et al., 2018; Ohta & Flajnik, 2006; Rogers et al., 2006). Knowledge at the genetic level of IgD in species other than human and rodents has only begun to accumulate recently, and has demonstrated that IGHD is widespread throughout vertebrates and is extremely diverse (Rogers et al., 2006). Homologs of IGHD ancestral genes can also be identified in amphibians and reptiles (Estevez et al., 2016; Gambón-Deza & Espinel, 2008), further underscoring its ancient evolutionary origins.

When compared with the sequences described in humans by Mills et al. (1990), the orangutan switch regions show strikingly similar features. In both species, Sμ and Sα regions are enriched in tandem pentamer repeats and display high GC content, whereas Sγ and Sε regions contain shorter, more fragmented repeats with lower motif density. The absence of a distinct switch-like region upstream of IGHD is consistent between orangutan and human, and the orangutan IGHGP likewise lacks canonical switch motifs. Thus, while the exact repeat structure and motif composition may differ, the relative distribution of strong and weak switch regions across the constant gene cluster is comparable between orangutan and human. In most mammals, expression of IGHD is generated primarily through alternative RNA splicing of a shared IGHM–IGHD primary transcript, rather than by classical class-switch recombination (CSR) (Arpin et al., 1998). Consequently, the switch region (Sδ) upstream of IGHD is generally weak or poorly defined in primates compared to the canonical Sμ, Sγ, Sα, and Sε elements (Chen et al., 2009; Watson and Breden, 2012). The absence of a strong Sδ region in orangutan assemblies is therefore consistent with the broader mammalian pattern, where CSR to IGHD is rare or nonfunctional. However, in *Pongo*, the IGHD gene itself is pseudogenized in all haplotypes examined, containing multiple disruptive mutations and frame shifts that likely abolish expression of a functional δ constant region. This distinguishes orangutans from humans and other apes, where IGHD remains intact and IGHD is expressed in naïve B cells (Dirks et al., 2023; Rogers et al., 2006). The combined absence of a canonical switch region and the pseudogenized IGHD sequence together suggest that functional IGHD expression may be lost in orangutans, representing a lineage-specific degeneration of this immunoglobulin isotype.

Previous studies have shown that IGHG copy number is highly variable across primates, particularly between New World monkeys and apes, indicating that IgG evolution has proceeded in lineage-specific ways. Within Hominidae, both gene gains and losses were common, and the resulting IGHG repertoires differ notably among species (Garzón-Ospina & Buitrago, 2022). In *Pongo* species, we annotated 6 IGHG genes: IGHG3A, IGHG3B, IGHG1, IGHG4, IGHGP, and IGHG2. Among these, IGHG3A, IGHG1, IGHG4, and IGHGP appear to be functional, while IGHG3B and IGHG2 are open reading frames (ORFs). In humans, copy number variation (CNV) is documented at the IGHG locus, particularly affecting IGHG4, which can occur in more than one copy in certain haplotypes (e.g., IGHG4A / duplicated IGHG4). Therefore, the total number of IGHG genes in humans is not fixed, and although the modal or reference haplotype is often described with five IGHG genes, some individuals may carry six IGHG genes including the duplicated IGHG4 (IGHG4A) (Papadaki et al., 2025). In Pongo, the coexistence of four functional IGHG genes together with two ORF copies is consistent with a birth-and-death mode of evolution, as described for primate IgG genes by Garzón-Ospina and Buitrago (2020). At the same time, the retention of orthologs shared with other great apes indicates that divergent evolution also contributes to the maintenance and functional specialization of conserved IGHG lineages.

The organization of the IGK and IGL loci in *Pongo abelii* and *Pongo pygmaeus* reveals a compact and functionally conserved architecture compared to the IGH locus. This reflects the well-documented evolutionary constraint imposed on light chain immunoglobulin genes across mammals. Consistent with observations in human, gorilla and rhesus macaque (Barbié & Lefranc, 1998; Debbagh et al., 2024; Nguefack Ngoune et al., 2022), both loci maintain stable genomic organization with minimal evidence of large-scale structural remodeling. This structural stability suggests that diversification in light chain loci occurs predominantly through allelic variation rather than alterations in gene copy number. Unlike humans, in whom the IGKV locus is divided into distinct proximal and distal clusters, orangutans show an organization more similar to that of gorillas, with a compact and continuous IGKV architecture.

RSS overall shows a high degree of sequence conservation in *Pongo* (Figure 5). With respect to the heptamer (canonical sequence: CACAGTG), the CAC sequence bordering the recombination site is perfectly conserved across all V genes of the *Pongo* locus. This conservation mirrors the pattern reported in humans and other mammals, where the CAC trinucleotide is maintained regardless of spacer length, associated coding sequence, or specific locus. Functional studies have shown that this trinucleotide is essential for efficient assembly of IG and TCR genes; mutations affecting any of the CAC positions markedly reduce RSS activity in recombination assays (Van Gent et al., 1995; Yin et al., 2009). The remaining four nucleotides of the heptamer motif are also well conserved among *Pongo* V RSSs, similar to the human pattern, indicating that strong evolutionary pressure maintains this motif to ensure proper recombination. In contrast, the nonamer displays more sequence variability among *Pongo* RSSs, particularly within the spacer-proximal nucleotides. Despite this heterogeneity, the nonamer positions 5-9 (AAACC) are well conserved in nearly all *Pongo* sequences, underscoring their critical functional role. This conservation asymmetry within the nonamer supports previous findings that highlight the importance of the AAA triplet in maintaining recombination efficiency. Variability in the first four nucleotides appears to have limited functional consequence, whereas changes affecting the conserved AAA region likely impair recombination activity. Overall, the *Pongo* RSS organization exhibits the same evolutionary logic observed in humans, maintaining the structural features essential for efficient V(D)J recombination (Akamatsu et al., 1994; Hesse et al., 1989; Ramsden et al., 1994).

## Conclusion

In this study, we present the first comprehensive genomic annotation of the immunoglobulin loci (IGH, IGK, and IGL) in Sumatran (*Pongo abelii*) and Bornean (*Pongo pygmaeus*) orangutans, analyzed across multiple high-quality genome assemblies. Through the application of the IMGT-ONTOLOGY and its rigorous curation framework, we ensured accurate gene identification, functionality assessment, and full alignment with IMGT nomenclature standards. Despite being based on a limited number of individuals, our analysis reveals a largely conserved IG locus organization with limited structural divergence between the two orangutan species. This characterization of IG locus diversity not only enhances our understanding of immune variation in great apes but also contributes valuable additions to the IMGT reference directories, thereby establishing a strong foundation for future investigations into antibody gene evolution, diversity, and function within the Hominidae family.

## Author Contributions

SB: Conceptualization, Data curation, Formal analysis, Investigation, Methodology, Resources, Validation, Visualization, Writing – original draft, Writing – review & editing. GZ: Assisted with the assembly and gene validation procedures, Writing – review & editing. CD: Conceptualization, early stage methodology training. GF, JJ-M and VG: Conceptualization, Data curation, Formal analysis, Investigation, Methodology, Validation, Visualization, Writing – review & editing. SK: Conceptualization, Funding acquisition, Project administration, Resources, Supervision, Validation, Writing – review & editing.

## Acknowledgments

We sincerely thank the IMGT® team for their dedication and steadfast enthusiasm, with special appreciation to Myriam Croze for her kindness, continuous support, and inspiring collegiality. We also acknowledge the internship students Julia Hensel, Vasiliki Zorlou and Leda Pothitaki who participated in IMGT biocuration training, typically for periods of two months.

## Supplementary data

The Supplementary Material for this article can be found online at: Supplementary_Notes_BATOOL

