## supplementary notes for "Characterization of Immunoglobulin Loci in *Pongo abelii* and *Pongo pygmaeus*: Insights from Multi-Genome Annotation and IMGT-Based Curation"

### 1. IMGT developed quality Control process

To investigate the immunoglobulin (IG) loci in orangutans, we analyzed all available genome assemblies (n=13) as of January 2024 from the NCBI Genome database. These assemblies represented two orangutan species (*Pongo abelii* & *Pongo pygmaeus*) and 7 assemblies were selected using IMGT's predefined two-phase pre-selection criteria available at https://www.imgt.org/IMGTScientificChart/Assemblies/IMGTassemblyquality.php. Assemblies were evaluated based on recency, assembly level, locus completeness, and sequencing/assembly methodology (Table S1).

| **Assembly name** | **Assembly ID/Accession** | **Species** | **Assembly Release date** | **Isolate** | **Sequencing Technology** | **Assembly Level** | **Used in Study** | **Comments** |
|---|---|---|---|---|---|---|---|---|
| NHGRI_mPonPyg2-v2.1_pri | GCA_028885625.3 | *Pongo pygmaeus* | Jan, 2024 | AG05252 | PacBio Sequel; Oxford Nanopore PromethION | Chromosome | Fully analysed | High-quality haploid assembly with full locus coverage and chromosome-level resolution. Supplementary Figure 1 (A, C and E) |
| NHGRI_mPonAbe1-v2.1_pri | GCA_028885655.3 | *Pongo abelii* | Jan, 2024 | AG06213 | PacBio Sequel; Oxford Nanopore PromethION | Chromosome | Fully analysed | High-quality haploid assembly with full locus coverage and chromosome-level resolution. Supplementary Figure 1 (B, D and F) |
| NHGRI_mPonPyg2-v2.0_alt | GCA_028885525.2 | *Pongo pygmaeus* | Jan, 2024 | AG05252 | PacBio Sequel; Oxford Nanopore PromethION | Chromosome | Fully analysed | High-quality haploid assembly with full locus coverage and chromosome-level resolution. Supplementary Figure 1 (A, C and E) |
| NHGRI_mPonAbe1-v2.0_alt | GCA_028885685.2 | *Pongo abelii* | Jan, 2024 | AG06213 | PacBio Sequel; Oxford Nanopore PromethION | Chromosome | Fully analysed | High-quality haploid assembly with full locus coverage and chromosome-level resolution. |

| | | | | | | | | |
|---|---|---|---|---|---|---|---|---|
| | | | | | | | | Supplementary Figure 1 (B, D and F) |
| Susie_PAB_hifiasm-v0.15.2.pri | GCA_030170355.1 | *Pongo abelii* | Jun, 2023 | Susie | PacBio Sequel | Contig | Used to fill gaps | Recent, high quality, full locus coverage, but on contigs. Was therefore used complementarily, to verify and fill sequence gaps within the reference locus and to confirm ambiguous regions Supplementary Figure 2 (A, C and E) |
| Susie_PAB_hifiasm-v0.15.2.alt | GCA_030170345.1 | *Pongo abelii* | Jun, 2023 | Susie | PacBio Sequel | Contig | Used to fill gaps | Recent, high quality, full locus coverage, but on contigs. Was therefore used complementarily, to verify and fill sequence gaps within the reference locus and to confirm ambiguous regions Supplementary Figure 2 (A, C and E) |
| mPonPyg1.1 | GCA_947095605.1 | *Pongo pygmaeus* | Oct, 2022 | | ONT | Contig | Not used | Partial, No locus coverage |
| ASM2376777v1 | GCA_023767775.1 | *Pongo pygmaeus* | Jun, 2022 | KIZ-2021_21 | PacBio RSII | Scaffold | Not used | Locus fragmented across multiple scaffolds |
| ponAbe.msY.makovalab.ver3 | GCA_015021835.1 | *Pongo abelii* | Oct, 2020 | 19910051_AG06213-MT | Illumina HiSeq | Scaffold | Not used | No locus coverage |
| Susie_PABv2 | GCA_002880775.3 | *Pongo abelii* | Jan, 2018 | Susie | PacBio; Illumina NextSeq 500; BioNano Saphyr (two enzyme) | Chromosome | Fully analysed | Good haploid representation and locus structure. Serves as reference in IMGT databases for *Pongo abelii* Supplementary Figure 2 (B, D and F) |

| | | | | | | | | |
|---|---|---|---|---|---|---|---|---|
| NGS12 | GCA_900086635.1 | *Pongo pygmaeus* | Jul, 2016 | | NGS Illumina HiSeq 2500 | Chromosome | Not used | Partial, No locus coverage |
| P_pygmaeus_2.0.2 | GCA_000001545.3 | *Pongo abelii* | Nov, 2008 | ISIS 71 | | Chromosome | Not used | No locus coverage |
| ASM16753v2 | GCA_000167535.2 | *Pongo abelii* | Nov, 2007 | | | Contig | Not used | Partial, No locus coverage |

Supplementary Table 1. **Assemblies assessed for immunoglobulin (IG) locus completeness and quality.** Thirteen genome assemblies were assessed for completeness and sequence quality of the immunoglobulin (IG) loci. Seven assemblies were retained (five fully analysed and two used for gap filling), whereas the others were excluded due to fragmentation or incomplete or missing IG locus coverage.

### 2. IMGT/StatAssembly

Inspired by Zhu et al. 's CloseRead tool (Zhu et al., 2025), IMGT/StatAssembly (version v0.1.4) (Zeitoun et al., 2025) is designed to analyze alignment data in BAM format. IMGT/StatAssembly uses as input the genomic positions of the IG and/or TR loci. For each specified IG/TR locus, the tool examines the region and counts the number of reads according to their mapping score. Any region covered by fewer than three primary reads is annotated as a “break” position. Secondary alignments (lower-quality mappings caused by repeats, as defined in minimap2 (Li, 2021)) and supplementary alignments (additional mappings from chimeric or split reads, also per minimap2 (Li, 2021)) are displayed together with the number of overlapping primary reads. High-quality assemblies are expected to show stable coverage, dominated by high mapping quality reads, with very few secondary, supplementary, or low-quality primary reads.

All assemblies used for analyzing IG loci in *Pongo abelii* and *Pongo pygmaeus* were evaluated using the IMGT/StatAssembly tool, preferably with PacBio HiFi reads, or PacBio SMRT reads in the case of the Susie PABv2 assembly, to generate BAM alignments. Reads were obtained from the publicly available NCBI BioProjects PRJNA916742, PRJNA916743, and PRJNA369439. The overall locus quality, as well as the quality of individual genes and alleles, was assessed based on the number of aligned primary reads. For each gene locus, the corresponding positions were provided to evaluate gene and allele integrity. An allele was considered confident if it was supported by the highest number of perfectly matching reads relative to the total read count.

In summary, IMGT/StatAssembly offers a robust approach for evaluating genome assembly quality directly from read data. It allows users to define custom threshold parameters to tailor the analysis to their needs. The tool also supports assessment of allele confidence by examining read support. When used alongside the established IMGT assembly quality criteria (Supplementary Table 1), it enables a systematic evaluation of both the overall assembly quality and the reliability of the resulting alleles. This methodology has been formally integrated into the standardized IMGT biocuration workflow and IMGT nomenclature procedures (https://www.imgt.org/IMGTScientificChart/Nomenclature/IMGTnomenclature.php).

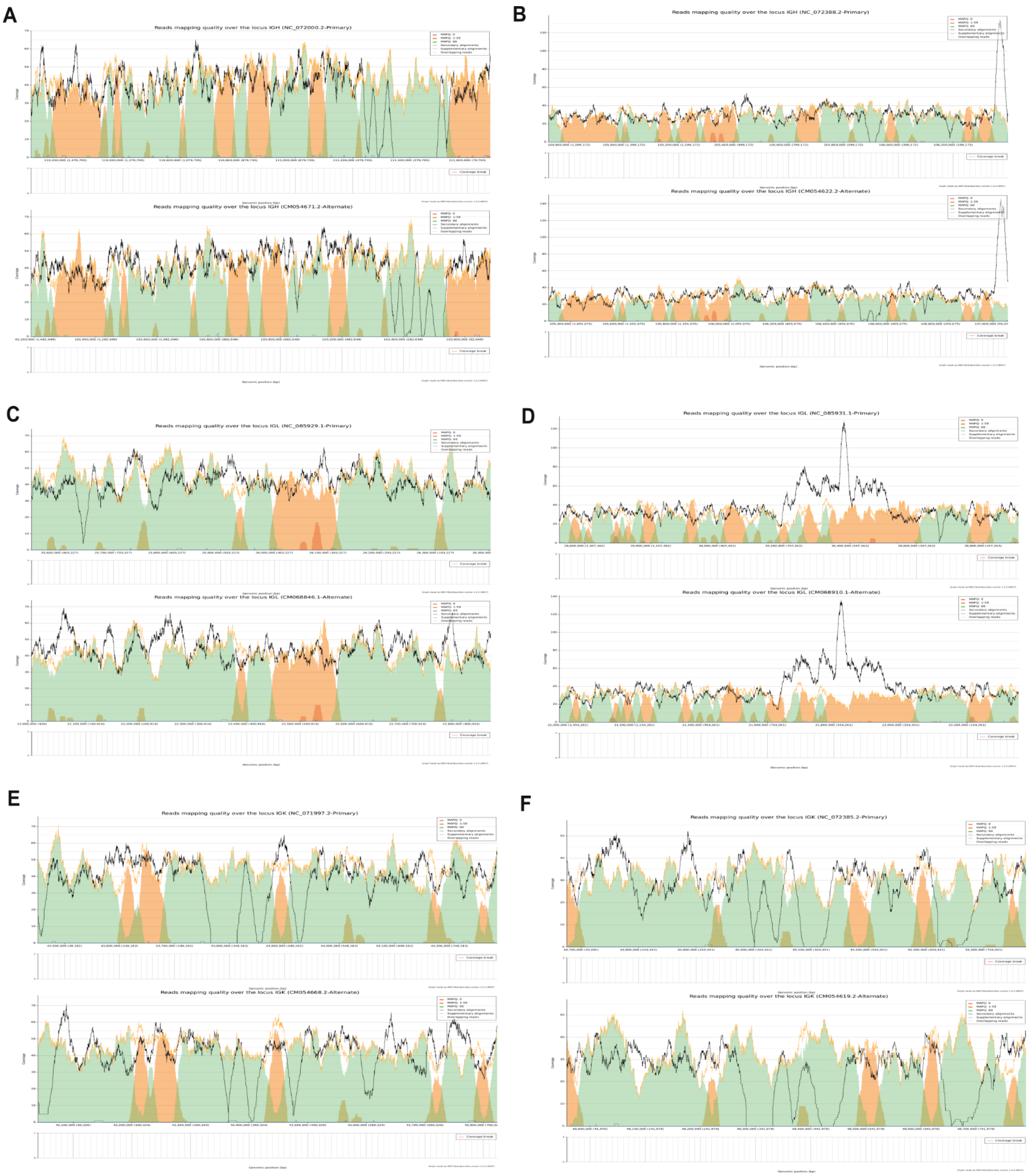

Supplementary Figure 1: **Locus quality assessment maps for orangutan immunoglobulin loci using IMGT/StatAssembly.** Panels A–F show locus-level read mapping quality across the immunoglobulin heavy (IGH), lambda (IGL), and kappa (IGK) loci for two *Pongo* species (left column = *Pongo abelii*; right column = *Pongo pygmaeus*), each represented by primary and alternate haplotype assemblies from NCBI. Each plot summarizes mapping quality profiles generated using IMGT/StatAssembly. Shaded areas indicate mapping quality categories (green, MAPQ = 60; orange, $1 \leq \text{MAPQ} < 60$; pink, MAPQ = 0), while the black and blue lines denote secondary and supplementary alignments across the locus. The x-axis corresponds to genomic position within each locus and the y-axis represents the locus coverage Chromosome accession numbers are noted in the title.

**(A)** IGH, *Pongo abelii* (NC_072000.2 / CM054671.2)
Both haplotypes show long continuous stretches of green (MAPQ 60), indicating broadly consistent, high-confidence read placement. Short orange segments (MAPQ 1–59) appear intermittently, mainly within internal regions, suggesting moderate mapping variability. Red intervals (MAPQ 0) are rare and short. Alignment traces (black/orange) are light and evenly distributed, showing that multi-mapping and supplementary reads are limited. The overall pattern is stable and uniform across the locus.
**(B)** IGH, *Pongo pygmaeus* (NC_072388.2 / CM054622.2)
Green regions dominate most of the locus in both haplotypes, reflecting generally high mapping confidence. Toward one end, a broader zone of orange and red occurs, together with denser alignment traces, indicating a localized area with lower mapping quality and higher read redundancy. Outside this region, color continuity and alignment density are similar to panel (A), showing overall comparable quality with a single distinct lower-MAPQ segment.
**(C)** IGL, *P. abelii* (NC_085929.1 / CM068846.1)
The color field is predominantly green with brief orange sections and few red intervals, showing consistently strong mapping quality. Alignment traces are thin and regular across the locus, and no extended low-MAPQ zones are present. The two haplotypes display nearly identical distributions, suggesting consistent assembly and read alignment behavior across both.
**(D)** IGL, *P. pygmaeus* (NC_085931.1 / CM068910.1)
A central region in both haplotypes contains an increase in orange and red bands and a higher density of alignment traces compared with flanking regions. The surrounding areas remain mostly green, showing normal mapping quality. The color contrast between the center and edges indicates one distinct segment with lower alignment confidence relative to the rest of the locus.
**(E)** IGK, *P. abelii* (NC_071997.2 / CM054668.2)
The IGK locus shows a repeating pattern of alternating green and orange bands extending along both haplotypes, reflecting a regular shift between high and moderate mapping quality across the locus. Red intervals are short and scattered. Alignment traces mirror the color alternation, appearing rhythmically distributed, with no extended blank or colorless gaps, showing that the mapping pattern is continuous and periodic.
**(F)** IGK, *P. pygmaeus* (NC_072385.2 / CM054619.2)
Both haplotypes display a color distribution almost identical to panel (E): alternating green and orange with rare red points and uniform alignment-trace density. The pattern repeats consistently along the entire locus, and the two species exhibit nearly the same mapping structure, indicating a comparable organization and data quality for IGK.

**Summary**

Across all loci, green predominates, showing that most regions have stable, high-confidence mappings. Orange and red intervals are restricted to specific, recurring positions, marking local zones of lower mapping certainty. Among loci, IGH and IGL vary more between species, while IGK patterns are highly consistent across both orangutans and between haplotypes. The color and alignment-trace patterns together indicate continuous, well-aligned loci with localized variability limited to discrete segments.

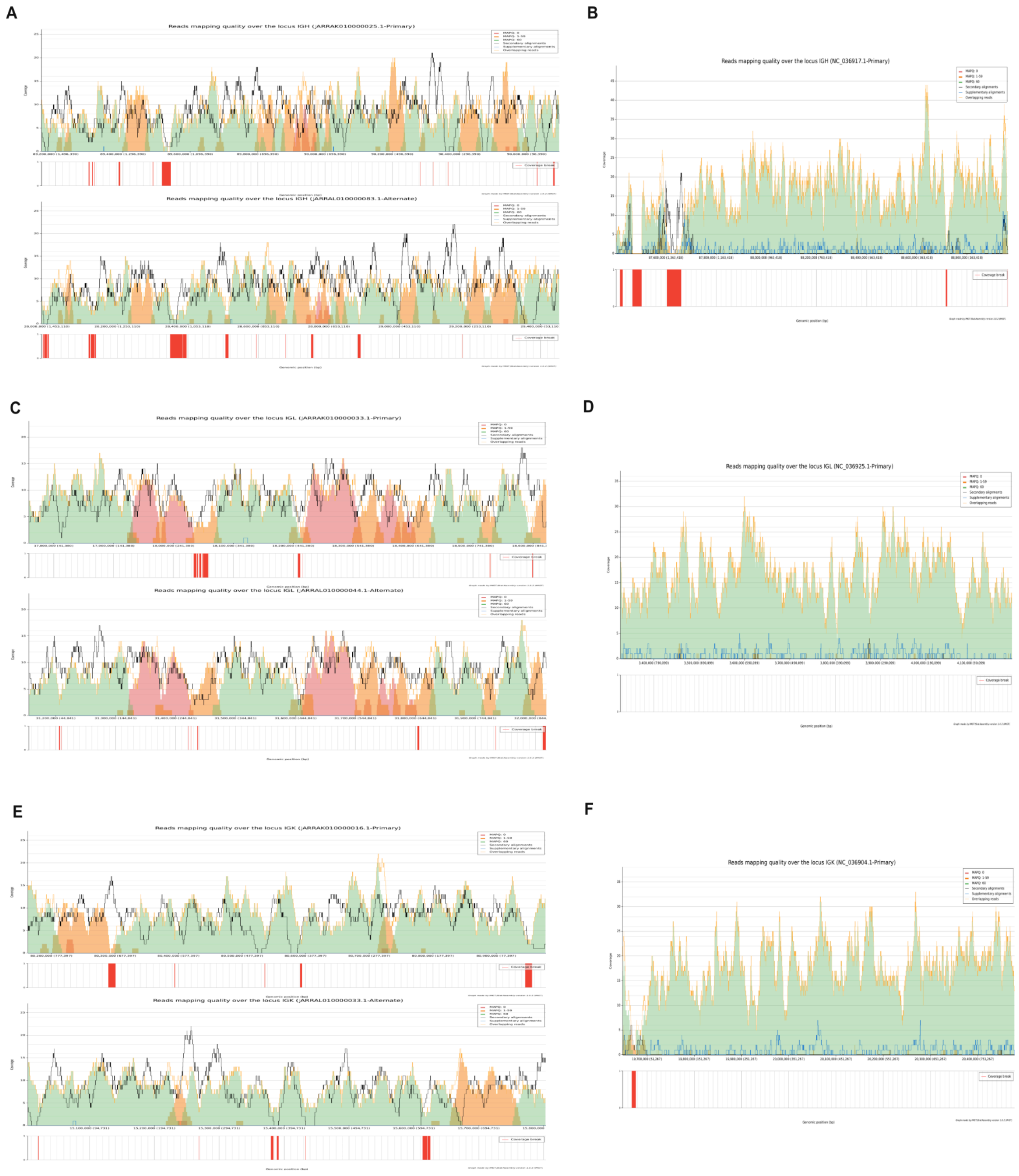

Supplementary Figure 2: **Locus quality assessment maps for *Pongo abelii* (Susie) immunoglobulin loci using IMGT/StatAssembly.** Panels A–F show locus-level read mapping quality across the immunoglobulin heavy (IGH), lambda (IGL), and kappa (IGK) loci for *Pongo* abelii (left column = Susie_PAB_hifiasm-v0.15.2.pri/alt; right column = Susie_PABv2). Each plot summarizes mapping quality profiles generated using IMGT/StatAssembly. Shaded areas indicate mapping quality categories (green, MAPQ = 60; orange, 1 ≤ MAPQ < 60; pink, MAPQ = 0), while the black and blue lines denote secondary and supplementary alignments across the locus. The x-axis corresponds to genomic position within each locus and the y-axis represents the locus coverage. Chromosome accession numbers are noted in the title.

**(A)** IGH (hifiasm pri/alt)
Both the primary and alternate haplotypes show extensive irregularity in read mapping quality across the IGH locus. Secondary alignments are frequent and widespread, confirming that many reads map equally well to multiple positions, while supplementary alignments mark boundaries between near-identical repeats. Coverage oscillates sharply between ~5× and >25×, and several no-coverage breaks are present, reflecting missing or unresolved sequences.
**(B)** IGH (Susie_PABv2)
In the Susie_PABv2 assembly, mapping quality is generally high (predominantly MAPQ 60) across the IGH locus, but several no-coverage breaks persist, particularly at the 5' end of the locus. Despite these localized breaks, the overall locus structure is more contiguous and mappability markedly higher than in the hifiasm assembly.
**(C)** IGL (hifiasm pri/alt)
In the Susie_PAB_hifiasm-v0.15.2 assembly, the IGL locus displays nearly identical mapping patterns between the primary and alternate haplotypes. Both show the same alternating bands of high (MAPQ 60) and low (MAPQ 0–59) mapping quality across the locus, along with matching peaks of secondary alignments. Coverage is uneven but closely mirrored between haplotypes, with recurring dropouts. Consequently, while the overall locus structure is captured, both haplotypes suffer from the same mapping ambiguity.
**(D)** IGL (Susie_PABv2)
In the Susie_PABv2 assembly, the IGL locus exhibits clean, uniform mapping with nearly all reads assigned a MAPQ of 60, indicating unique alignment throughout the region. Secondary and supplementary alignments are minimal, and no coverage breaks are detected, suggesting that the locus is represented as a single, contiguous sequence without apparent collapse or structural gaps.
**(E)** IGK (hifiasm pri/alt)
In the Susie_PAB_hifiasm-v0.15.2 assembly, the IGK locus shows extensive mapping ambiguity in both the primary and alternate haplotypes. The alternate contig closely resembles the primary, sharing nearly identical regions of low mapping quality and coverage breaks
**(F)** IGK (Susie_PABv2)
In the Susie_PABv2 assembly, the IGK locus shows consistently high mapping quality, with nearly all reads aligning uniquely (MAPQ 60) across the region. Secondary and supplementary alignments are rare and of low magnitude. Coverage is smooth and continuous, with only a single localized dropout near the 3' end of the locus.

**Summary**

Although the Susie_PABv2 assembly exhibits higher mapping quality and greater sequence contiguity overall, it still contains several short no-coverage gaps, particularly within the IGH and IGK loci. The hifiasm contigs often span regions that are missing or weakly represented in PABv2, while PABv2 resolves areas that remain ambiguous in the HiFi assembly. This complementarity indicates that combining information from both assemblies would provide a more complete and contiguous reconstruction of the IGH, IGK, and IGL loci in Susie.

#### 3. Genetic Structure of IG loci

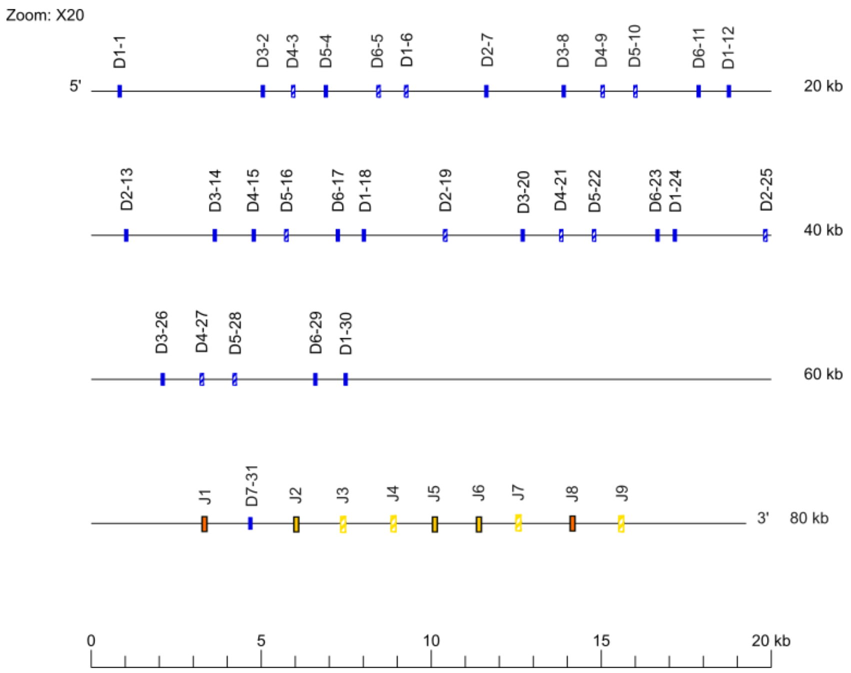


Supplementary Figure 3: 20x Zoom view of D-J-CLUSTER in IGH locus.

| *Pongo* Haplotypes | IGHV | | | IGHD | | | IGHJ | | | IGHC | | | Total |
|---|---|---|---|---|---|---|---|---|---|---|---|---|---|
| | P | ORF | F | P | ORF | F | P | ORF | F | P | ORF | F | |
| Susie | 87 | 5 | 47 | 0 | 12 | 19 | 2 | 4 | 3 | 5 | 1 | 7 | 192 |
| AG06213 hap1_alt | 91 | 5 | 44 | 0 | 12 | 19 | 2 | 4 | 3 | 1 | 2 | 9 | 192 |
| AG06213 hap2_pri | 100 | 7 | 51 | 0 | 12 | 19 | 2 | 4 | 3 | 1 | 2 | 9 | 210 |
| AG05252 hap1_alt | 110 | 7 | 59 | 0 | 12 | 19 | 2 | 4 | 3 | 1 | 2 | 9 | 228 |
| AG05252 hap2_pri | 104 | 10 | 53 | 0 | 12 | 19 | 2 | 4 | 3 | 1 | 2 | 9 | 219 |

Supplementary Table 2: Distribution of functional, open reading frame, and pseudogene segments across IGHV, IGHD, IGHJ, and IGHC loci in five *Pongo* haplotypes.

| *Pongo* Haplotypes | IGLV | | | IGLJ | | | IGLC | | | Total |
|---|---|---|---|---|---|---|---|---|---|---|
| | P | ORF | F | P | ORF | F | P | ORF | F | |
| Susie | 72 | 3 | 30 | 0 | 3 | 5 | 2 | 0 | 5 | 120 |
| AG06213 hap2_alt | 77 | 2 | 25 | 0 | 3 | 5 | 2 | 0 | 5 | 119 |
| AG06213 hap1_pri | 71 | 3 | 29 | 0 | 3 | 5 | 2 | 0 | 5 | 118 |
| AG05252 hap1_alt | 79 | 0 | 34 | 0 | 2 | 5 | 2 | 0 | 5 | 127 |
| AG05252 hap2_pri | 77 | 1 | 36 | 0 | 2 | 5 | 2 | 0 | 5 | 128 |

Supplementary Table 3: Distribution of functional, open reading frame, and pseudogene segments across IGLV, IGLJ, and IGLC loci in five *Pongo* haplotypes.

| *Pongo* Haplotypes | IGKV | | | IGKJ | | | IGKC | | | Total |
|---|---|---|---|---|---|---|---|---|---|---|
| | P | ORF | F | P | ORF | F | P | ORF | F | |
| Susie | 42 | 5 | 30 | 1 | 0 | 4 | 0 | 0 | 1 | 83 |
| AG06213 hap2_alt | 39 | 4 | 30 | 1 | 0 | 4 | 0 | 0 | 1 | 79 |
| AG06213 hap1_pri | 40 | 4 | 31 | 1 | 0 | 4 | 0 | 0 | 1 | 81 |
| AG05252 hap1_alt | 36 | 4 | 33 | 1 | 0 | 4 | 0 | 0 | 1 | 79 |
| AG05252 hap2_pri | 38 | 5 | 34 | 0 | 0 | 5 | 0 | 0 | 1 | 83 |

Supplementary Table 4: Distribution of functional, open reading frame, and pseudogene segments across IGKV, IGKJ, and IGKC loci in five *Pongo* haplotypes.

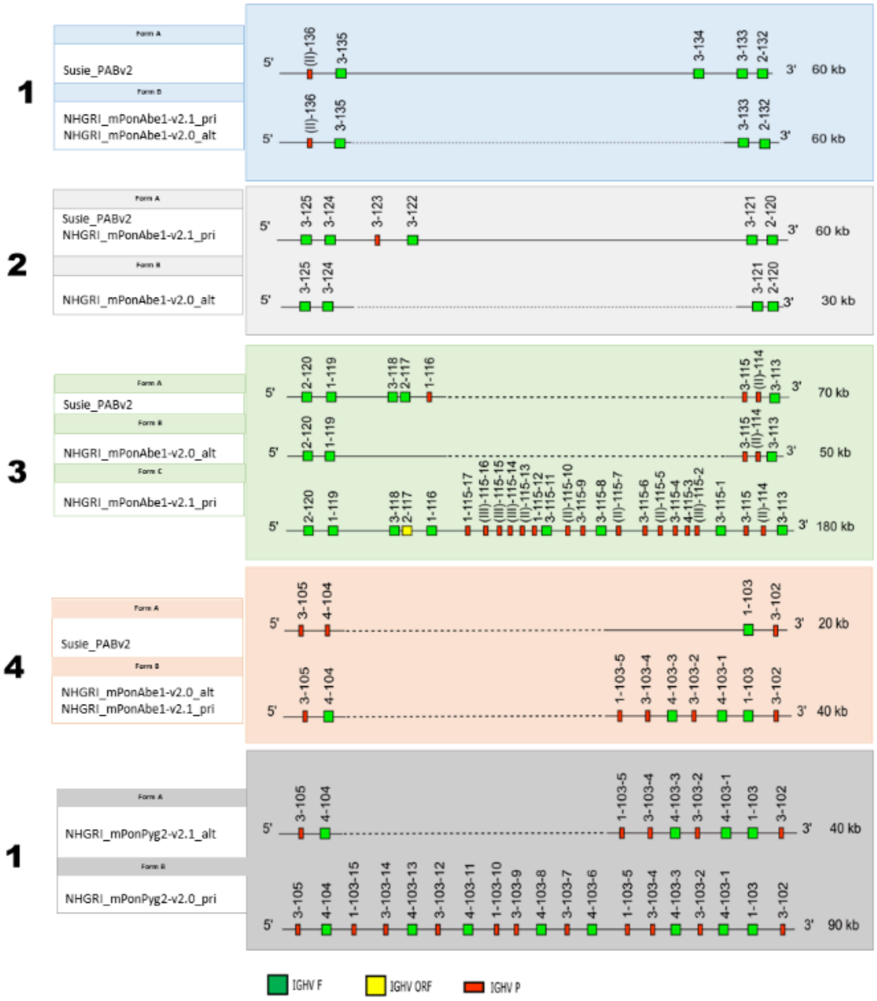


Supplementary Figure 4: Shaded regions, corresponding to Figure 1 in the main text, represent haplotype-specific insertions, deletions, or structural variants, validated by comparison across haplotypes and individuals from both Pongo abelii and Pongo pygmaeus, and further supported by ultra-long read alignments visualized in IGV.

### 4. Read-level Validation of Structural Variants Using IGV Visualization

To evaluate the structural correctness of IG loci across assemblies, we manually inspected each region in IGV (Integrative Genomics Viewer v2.18.5+dfsg-1) using Oxford Nanopore long-read alignments against the corresponding reference scaffolds. Reads were displayed with small indels masked (<20 bp) and alignment shading based on mapping quality, allowing us to distinguish true sequence absence from low-confidence or misassembled regions. We examined coverage continuity, soft-clipped or split-read signals, and changes in mapping quality to differentiate genuine gene absence from assembly collapse, as well as to detect false duplications or misassembled fragments.

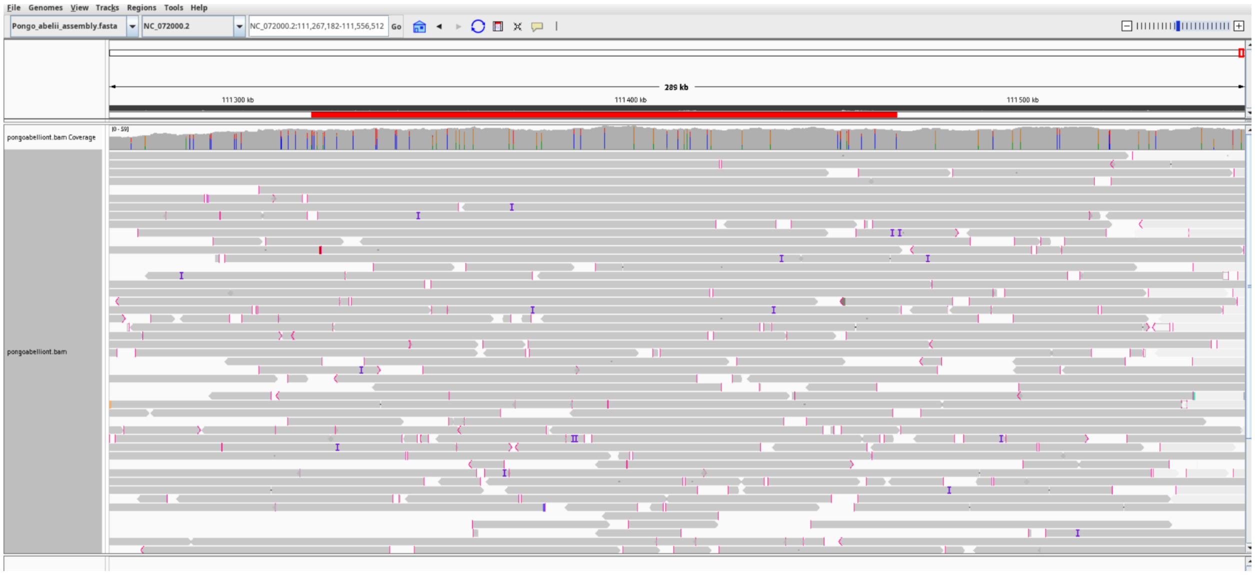


Supplementary Figure 5: **Read-based validation of the IGHV3-115–IGHV1-116 interval in *Pongo abelii*.** Oxford Nanopore long-read alignments over the IGHV3-115–IGHV1-116 region demonstrate continuous, high-quality mapping across both haplotypes. In the primary assembly, reads span a ~127 kb interval containing additional IGHV genes between IGHV3-115 and IGHV1-119, while in the alternate assembly, the two genes are directly adjacent with no intervening annotated segments. In both cases, reads align seamlessly without breaks or discordant mappings, confirming that each configuration represents a genuine haplotypic structure rather than an assembly error.

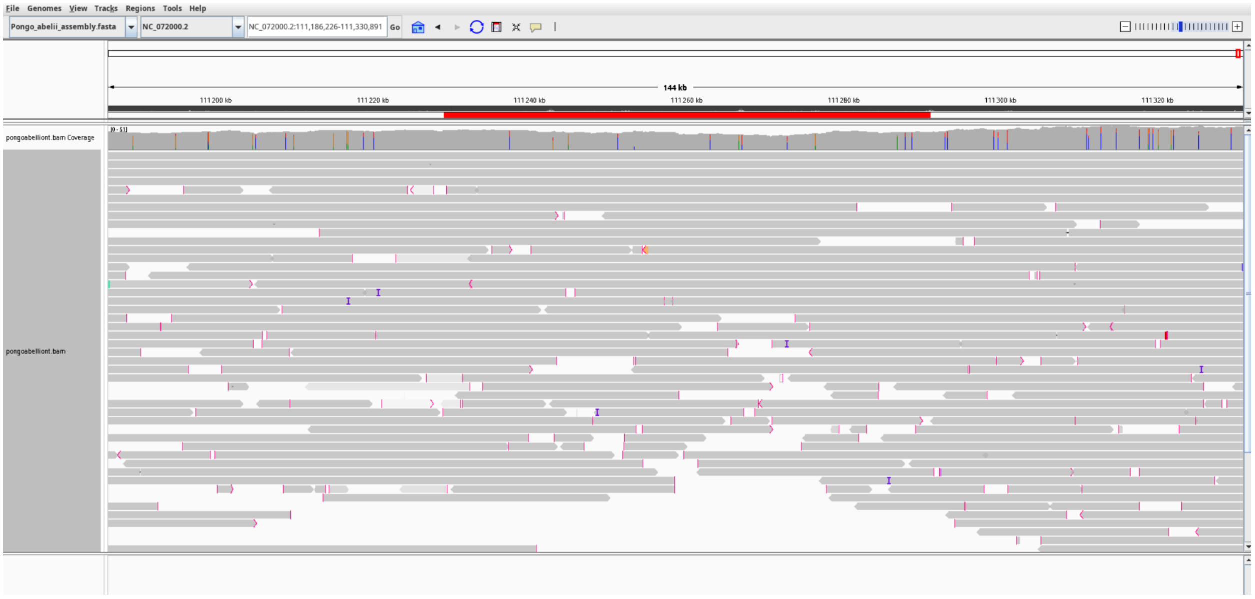

Supplementary Figure 6: **Read-based validation of the IGHV4-99–IGHV4-104 interval in *Pongo abelii*.** Oxford Nanopore long-read alignments from the NHGRI_mPonAbe1-v2.0 primary assembly show continuous, high-quality mapping across the 48.7 kb region spanning IGHV4-99 to IGHV4-104 (NC_072000.2:111,237,572–111,286,265). Reads align seamlessly without split mappings or coverage interruptions. The alternate haplotype displays the same contiguous read alignment pattern, indicating that this configuration is consistent and preserved across both haplotypes.

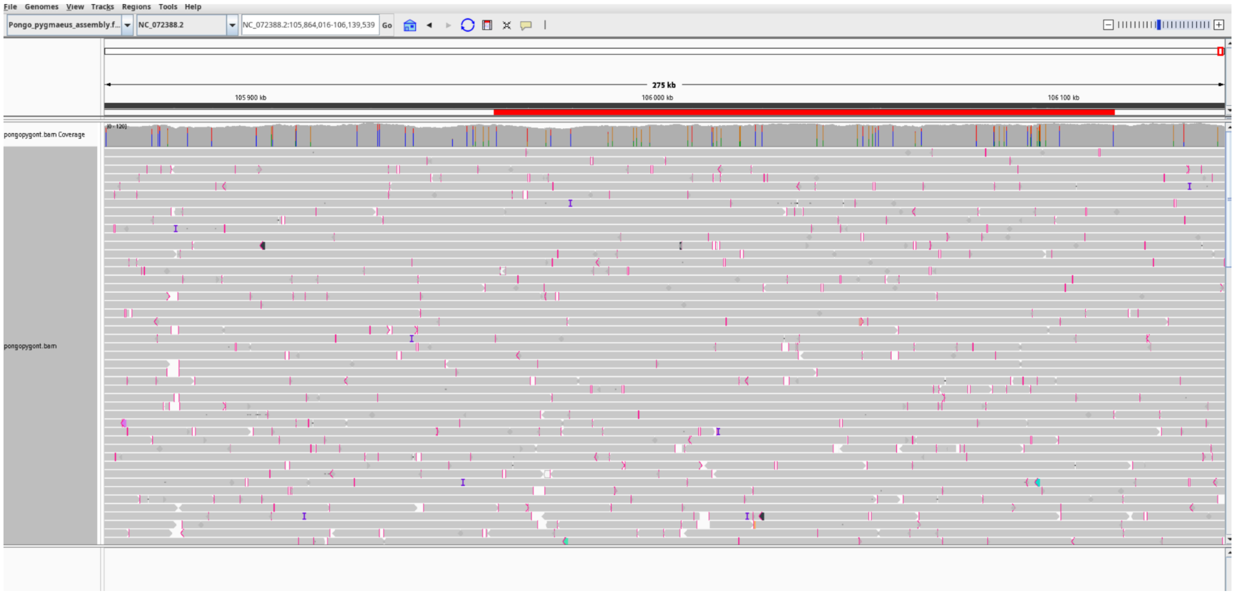

Supplementary Figure 7: **Read-based confirmation of the IGHV3-115–IGHV1-116 interval in Pongo pygmaeus.** Oxford Nanopore read alignments from the NHGRI_mPonPyg2-v2.0 primary assembly show continuous, high-quality coverage across the 127 kb region spanning IGHV3-115 to IGHV1-116 (NC_072388.2:105,977,156–106,104,187). Reads align seamlessly without split mappings or coverage gaps, confirming that the interval is structurally intact and authentically represented. The alternate haplotype exhibits the same uninterrupted alignment pattern, indicating that the IGHV3-115–IGHV1-116 configuration is consistent and preserved across both haplotypes

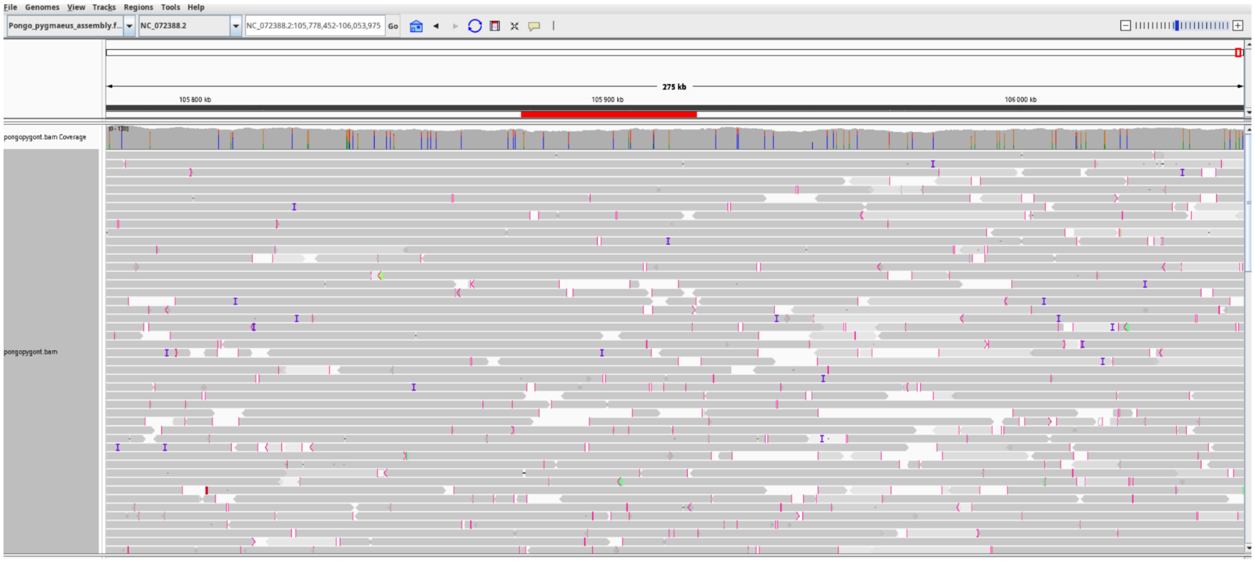


Supplementary Figure 8: **Read-based validation of the IGHV1-98–IGHV4-104 interval in *Pongo pygmaeus*.** Oxford Nanopore long-read alignments from the NHGRI_mPonPyg2-v2.0_pri assembly show continuous, high-quality mapping across the region spanning IGHV1-98 to IGHV4-104 (NC_072388.2:105895377-105924689). Reads align seamlessly without split mappings or coverage interruptions.

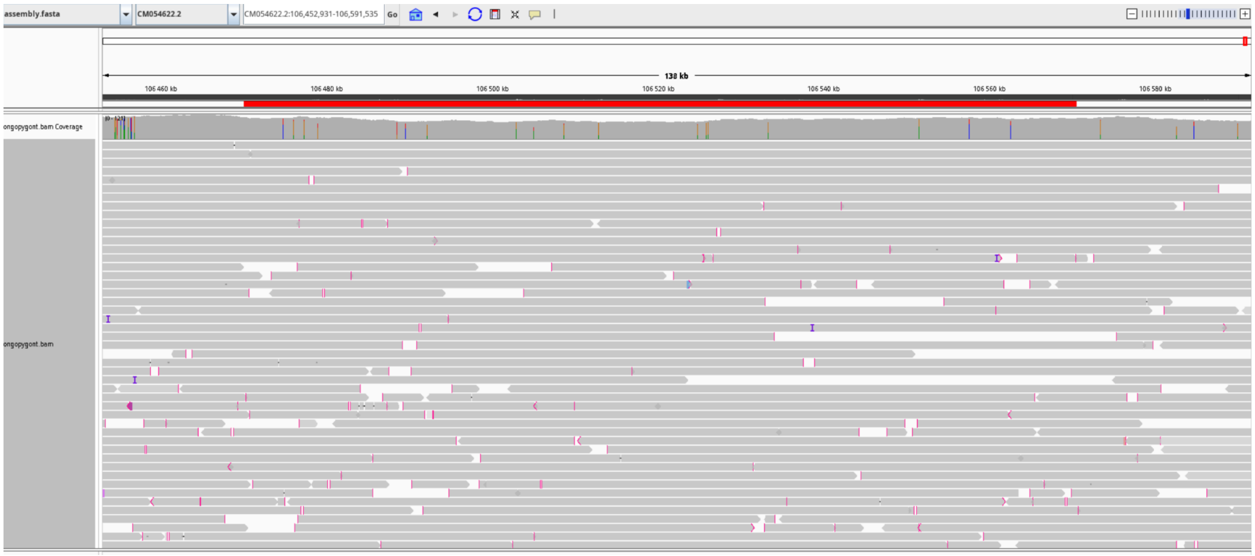


Supplementary Figure 9: **Read-based validation of the IGHV1-98–IGHV4-104 interval in *Pongo pygmaeus*.** Oxford Nanopore read alignments across the 138 kb region spanning IGHV1-98 to IGHV4-104 (CM054622.2:106,473,521–106,570,944) in NHGRI_mPonPyg2-v2.0_alt assembly show continuous, high-quality coverage with uniformly aligned reads and no coverage gaps or split mappings.

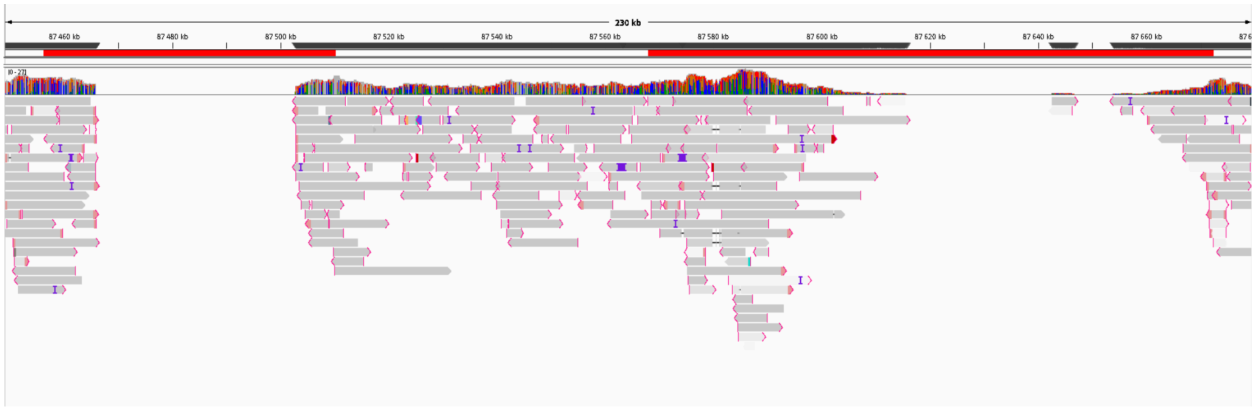


Supplementary Figure 10: **Read mapping evidence for IGHC region anomalies in Susie_PABv2.** Read alignments across the IGHC locus in the Susie_PABv2 assembly reveal two distinct regions of concern. The first corresponds to a complete coverage gap, where no reads align, matching the genomic position where IGHGP and IGHG2 should occur, indicating a local assembly collapse rather than true gene deletion. The second region showing irregular, fragmented read alignments at the annotated IGHGP1, IGHEP, and IGHG3C sites lacks continuous, high-quality read support, indicating that these genes are assembly artifacts rather than genuine genomic features.

### 5. Analysis of RSS

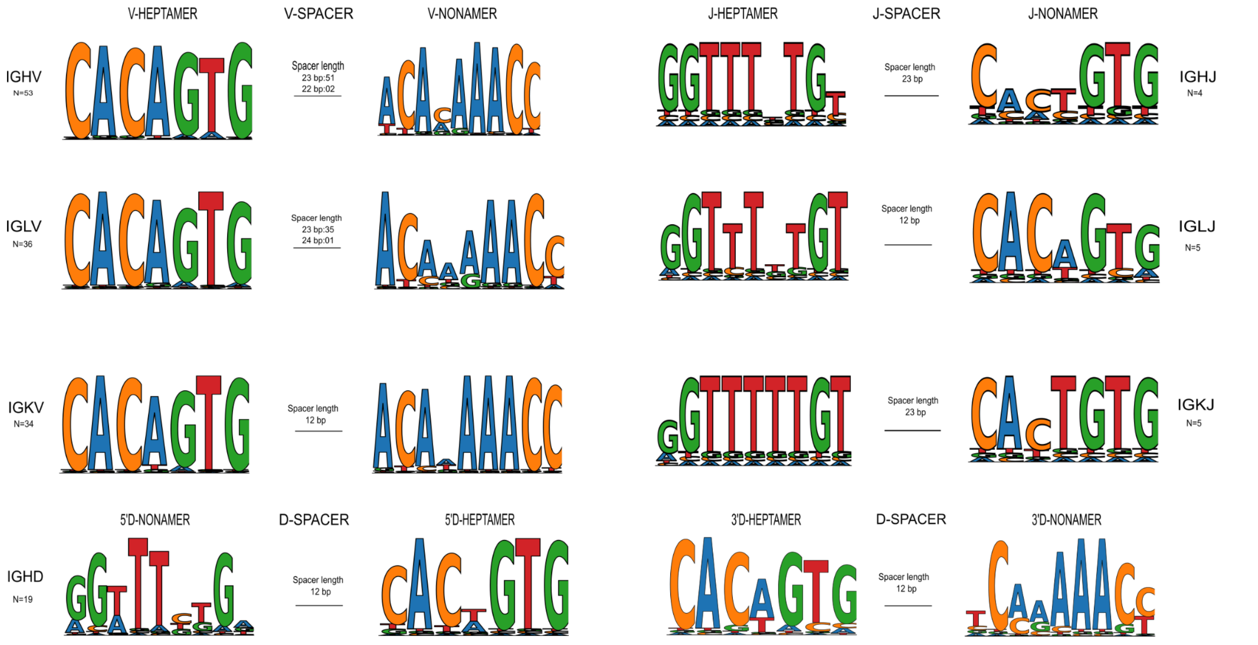


Supplementary Figure 11: **Consensus motifs of recombination signal sequences (RSSs) from functional immunoglobulin genes in *Pongo pygmaeus*.** Sequence logos derived from functional IG V, D, and J genes highlight the highly conserved heptamer and nonamer motifs, which flank coding segments and direct RAG-mediated recombination. Both motifs show strong positional conservation, particularly within the core CAC and ACA elements, consistent with their essential role in synapsis and cleavage. The observed spacer length distribution confirmed the expected 12/23-bp rule, with minor locus-specific sequence variability suggesting potential fine-tuning of recombination efficiency across loci.

### 6. Switch Region Analysis

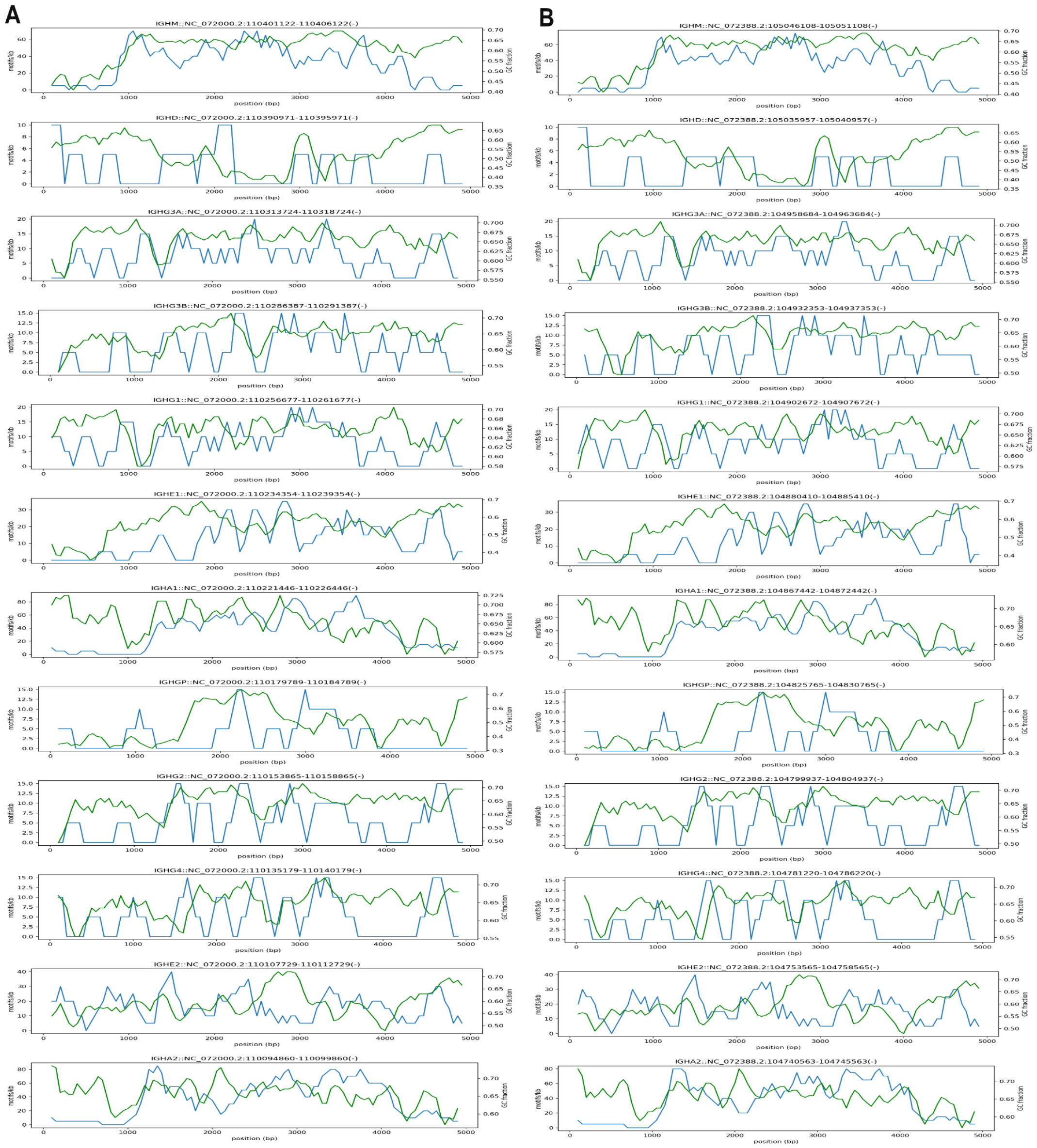


Supplementary Figure 12: Quality control (QC) plots showing sliding-window distributions of motif density (green) and GC fraction (blue) across immunoglobulin heavy chain (IGH) switch regions in *Pongo abelii* (A, left) and *Pongo pygmaeus* (D, right). Each subplot represents an individual switch (S) region located upstream of a constant region gene (e.g., Sμ, Sδ, Sγ, Sε, Sα). Peaks in motif density and GC fraction indicate the canonical G-rich, repetitive architecture characteristic of functional S regions.

Consistent compositional profiles across both species confirm accurate extraction and conserved sequence structure within the *Pongo* IGH constant locus.

| Gene | GAGCW_count | GAGCW_per_kb | GGGBT_count | GGGBT_per_kb | GC_fraction |
|---|---|---|---|---|---|
| IGHM | 175 | 35 | 240 | 48 | 0.61 |
| IGHD | 13 | 2.6 | 24 | 4.8 | 0.54 |
| IGHG3A | 92 | 18.4 | 36 | 7.2 | 0.659 |
| IGHG3B | 58 | 11.6 | 40 | 8 | 0.643 |
| IGHG1 | 105 | 21 | 42 | 8.4 | 0.659 |
| IGHE1 | 51 | 10.2 | 134 | 26.8 | 0.562 |
| IGHA1 | 170 | 34 | 251 | 50.2 | 0.657 |
| IGHGP | 9 | 1.8 | 21 | 4.2 | 0.491 |
| IGHG2 | 58 | 11.6 | 41 | 8.2 | 0.644 |
| IGHG4 | 52 | 10.4 | 38 | 7.6 | 0.65 |

Supplementary Table 5: **Motif frequencies and GC content in 5 kb upstream regions of *Pongo abelii* IGHC genes.** Counts of degenerate switch motifs were measured in the 5 kb upstream region of each constant region gene. GAGCW (where W = A or T) and GGGBT (where B = C, G, or T) represent canonical switch pentamers targeted by activation-induced cytidine deaminase. Values are reported as total motif counts and normalized density per kilobase (per_kb). GC_fraction denotes the proportion of G and C bases in each region. Strong enrichment of motifs and elevated GC fractions are evident upstream of IGHM and IGHA1/2, consistent with classical Sμ and Sα regions, whereas IGHD and IGHGP lack such features. Intermediate densities were observed for IGHG and IGHE genes.

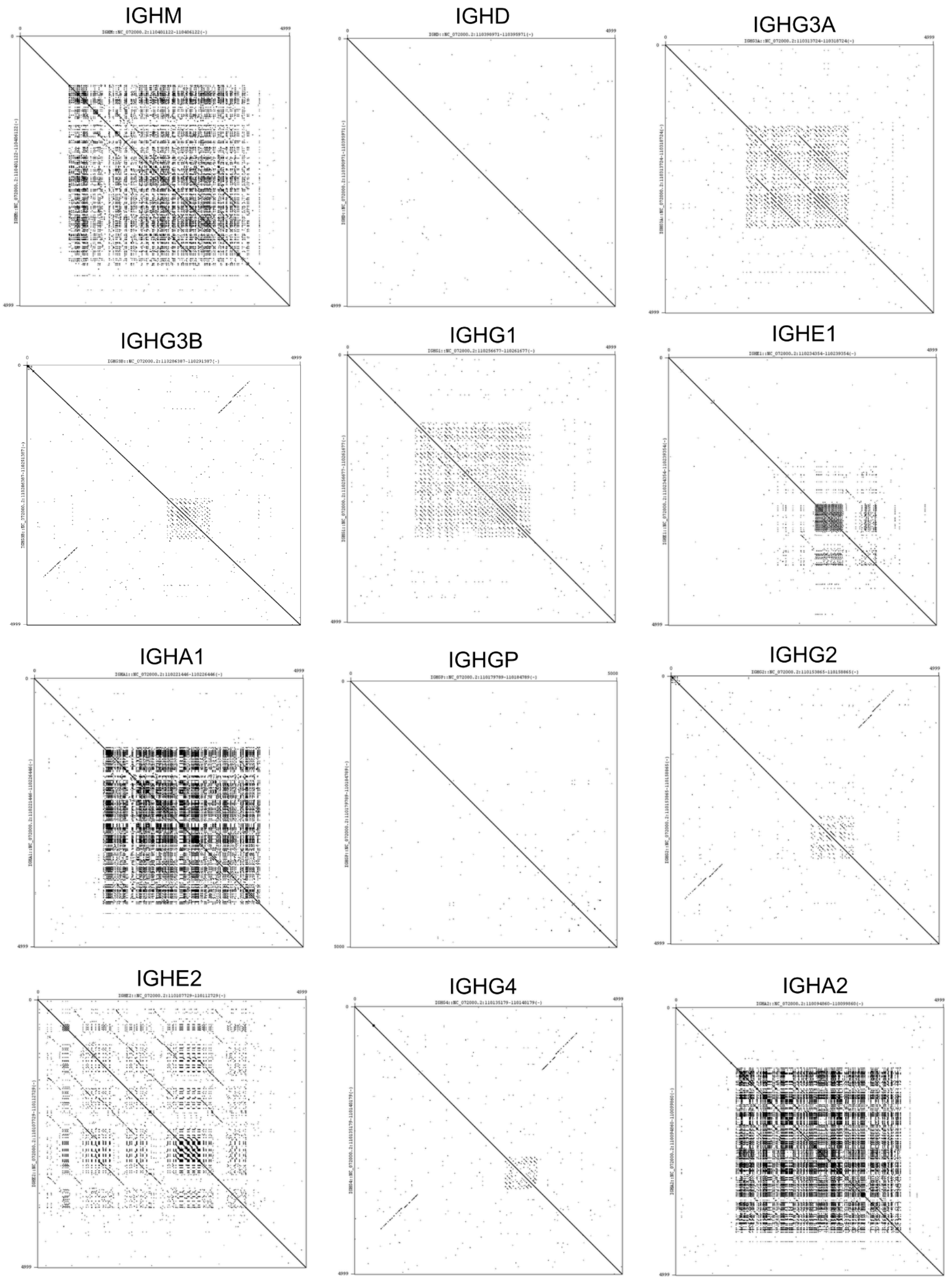


Supplementary Figure 13: **Dotplot of the IGHC genes in *Pongo abelii*.** Per-gene dotplots of 5 kb upstream regions validate the patterns observed at the locus level, with each constant region gene shown separately (word size 10).

### 7. Locus size vs Gene count

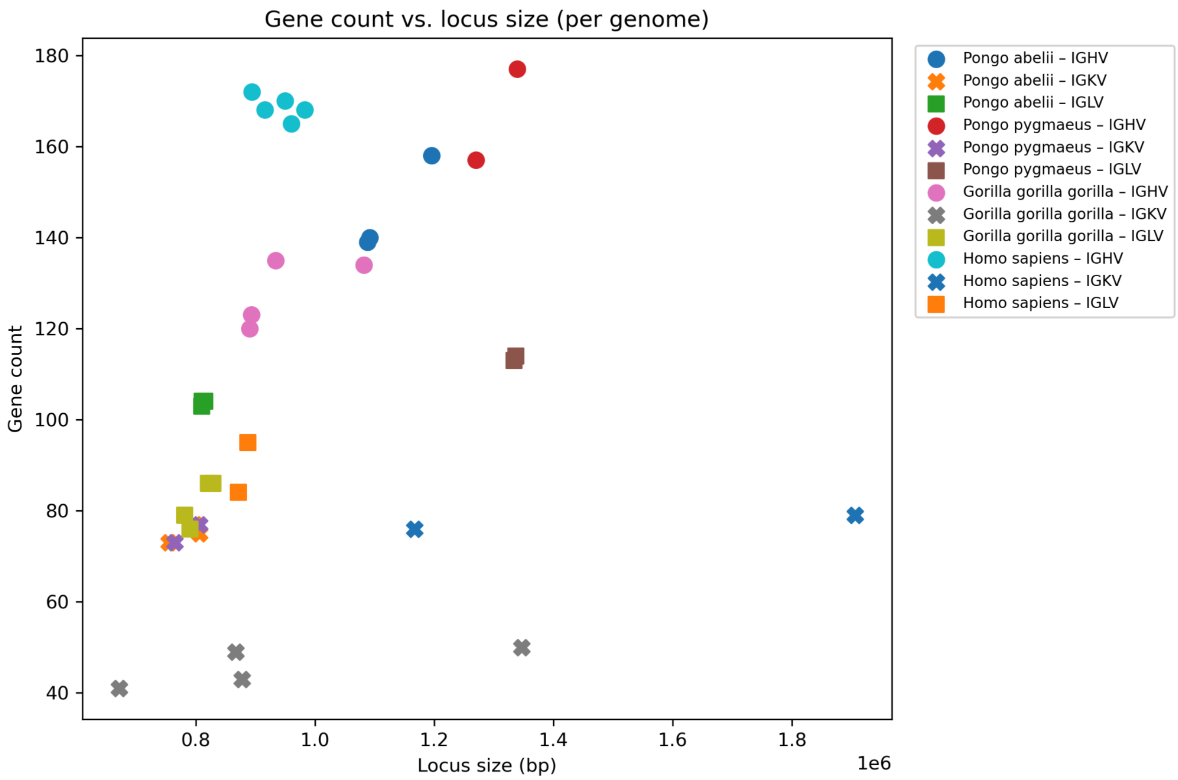


Supplementary Figure 14: **Relationship between immunoglobulin locus size and gene count across primate genomes.** Scatter plot showing the relationship between locus size (in base pairs) and total gene count for the immunoglobulin heavy (IGH), kappa (IGK), and lambda (IGL) loci across *Pongo abelii, Pongo pygmaeus, Gorilla gorilla gorilla,* and *Homo sapiens.* The correlation between locus length and gene number is strongest for the IGH locus, indicating that genomic expansion contributes to repertoire diversity in great apes.

| Species | Locus | Locus length | V-CLUSTER | Genes | IMGT Accession numbers | Locus | Locus length | V-CLUSTER | Genes | IMGT Accession numbers | Locus | Locus length | V-CLUSTER | Genes | IMGT Accession numbers |
|---|---|---|---|---|---|---|---|---|---|---|---|---|---|---|---|
| *Pongo abelii* | IGH | 1561328 bp | 1088121 | 139 | IMGT000122 | IGL | 863163 bp | 815129 | 104 | IMGT000108 | IGK | 828105 bp | 798671 | 77 | IMGT000096 |
| | | 1492363 bp | 1092016 | 140 | IMGT000121 | | 855536 bp | 810319 | 103 | IMGT000246 | | 834843 bp | 806000 | 75 | IMGT000244 |
| | | 1599061 bp | 1195740 | 158 | IMGT000206 | | 855847 bp | 810629 | 104 | IMGT000247 | | 783768 bp | 754919 | 73 | IMGT000245 |
| *Pongo pygmaeus* | IGH | 1745987 bp | 1339531 | 177 | IMGT000229 | IGL | 1378873 bp | 1333666 | 113 | IMGT000248 | IGK | 797359 bp | 764914 | 73 | IMGT000275 |
| | | 1672821 bp | 1270257 | 157 | IMGT000230 | | 1382051 bp | 1336848 | 114 | IMGT000278 | | 832478 bp | 806195 | 77 | IMGT000276 |
| *Gorilla gorilla gorilla* | IGH | 1141211 bp | 933797 | 135 | IMGT000074 | IGL | 838685 bp | 790466 | 76 | IMGT000086 | IGK | 892858 bp | 877825 | 43 | IMGT000077 |
| | | 1421867 bp | 1082038 | 134 | IMGT000078 | | 876832 bp | 828625 | 86 | IMGT000094 | | 698223 bp | 671684 | 41 | IMGT000089 |
| | | 1311160 bp | 890210 | 120 | IMGT000139 | | 826751 bp | 781363 | 79 | IMGT000152 | | 1373629 bp | 1346898 | 50 | IMGT000137 |
| | | 1315397 bp | 893859 | 123 | IMGT000140 | | 872234 bp | 821465 | 86 | IMGT000153 | | 893572 bp | 866928 | 49 | IMGT000138 |
| *Homo sapiens* | IGH | 1293408 bp | 950000 | 170 | IMGT000035 | IGL | 916838 bp | 871511 | 84 | IMGT000225 | IGK | 1944578 bp | 1905792 | 79 | IMGT000281 |
| | | 1331461 bp | 960793 | 165 | IMGT000110 | | 932853 bp | 887529 | 95 | IMGT000280 | | 1398008 bp | 1166956 | 76 | IMGT000100 |
| | | 1259983 bp | 915996 | 168 | IMGT000130 | | | | | | | | | | |
| | | 1249050 bp | 894048 | 172 | IMGT000113 | | | | | | | | | | |
| | | 1336686 bp | 983045 | 168 | IMGT000147 | | | | | | | | | | |

Supplementary Table 6. IMGT locus data underlying the V-gene density analysis (Figure 11 A), showing locus length, V-cluster size, and gene counts for IGH, IGL, and IGK loci across *Pongo abelii, Pongo pygmaeus, Gorilla gorilla gorilla and Homo sapiens* genomes.
